# Reversible decrease in swimming motility accompanies emergent β-lactam antibiotic resistance in *Escherichia coli*

**DOI:** 10.64898/2026.11.06.764950

**Authors:** Lisa M. Stabryla, William E. Ackerman, Jennifer N. Dootz, Stephanie L. Servetas, Scott A. Jackson, Jason G. Kralj

## Abstract

Antimicrobial resistance (AMR) threatens global health, food security, and development; however, the evolutionary and molecular pathways that drive AMR remain poorly understood, limiting our ability to advance diagnostics and rapidly detect newly emerging resistance. In this report, we employed laboratory evolution of an environmental *Escherichia coli* strain (NIST 0056) under β-lactam antibiotic pressure (ampicillin and cephalexin) to specifically investigate the relationship between resistance genotype and motility phenotype. Within 10 passages, populations evolved varying levels of β-lactam resistance (including high-level resistance > 32 µg/mL) and showed consistent, dose-dependent reductions in swimming motility. Whole-population genome sequencing revealed large deletions in *marR*, SNPs in efflux pump gene *acrB* and fumarate reductase gene *frdD,* and mutations with evidence of amplification in the chromosomal class C β-lactamase gene *ampC,* collectively linking these efflux-mediated, β-lactam resistance mechanisms to loss of motility. Notably, motility reductions were more pronounced in individual clones than in the mixed populations (*p* < 0.005, Mann-Whitney; two-tailed test). Following removal of antibiotic pressure, motility was restored and AMR-associated mutations decreased in frequency, suggesting fitness costs associated with resistance. In aggregate, our results demonstrate that reduced swimming motility accompanies specific efflux-driven, β-lactam resistance mechanisms. Further work will investigate if the behavior is generalizable across flagellated Enterobacteriaceae. Because these rapid changes are amenable to high-throughput measurement, swimming motility may offer a tractable physiological metric for AMR detection in environmental surveillance.

## 1. Introduction

Antimicrobial resistance (AMR) poses a profound threat to global public health and food security, and is considered one of the greatest systemic challenges of our time^1,2^. In response, targeted monitoring of AMR, including both wastewater and general environmental surveillance, has been identified as a critical strategy in combatting and managing resistance by the World Health Organization (WHO) and the U.S. Centers for Disease Control and Prevention (CDC).^1–8^

In line with this, development of rapid, high-throughput strategies is required to improve detection, identification, and characterization of AMR at scale. Indeed, the creation of such technologies was outlined as a major goal in the 2020-2025 National Action Plan for combating AMR.^3,5,6^ Detecting and monitoring AMR in human, agricultural, and environmental samples presents a formidable challenge owing to the diversity and spatiotemporal dynamism of microbial targets. Current AMR detection methods rely on culture-based strategies to assess antimicrobial susceptibility (*e.g.*, disk diffusion, antimicrobial gradient tests, broth microdilution, *etc.*); these are typically time- and cost-intensive (often requiring more than 24 h), with limited throughput and poor potential for automation.

On the other hand, emerging methods utilizing next-generation sequencing (NGS) technologies focus on underlying genetic and molecular information to predict AMR phenotypes. While these methods are high-throughput, they remain expensive and time-consuming. Additionally, NGS requires specialized training and may lack sensitivity (particularly when genotype-phenotype relationships are complex or insufficiently understood).^9–11^ Thus, there remains an unmet need for new, alternative, or complementary technologies that enable rapid, simple, and higher-throughput detection and identification of resistant bacteria. The development of such technologies will be useful for identifying AMR hotspots in the environment, informing intervention strategies and policy development, and assessing human health risk.

In clinical settings, antimicrobial susceptibility testing (AST), including MIC-based assays, remains the cornerstone for guiding treatment decisions, yet these measurements capture only a narrow snapshot of bacterial resistance. MIC values report growth inhibition under standardized conditions, but fail to account for the fitness consequences of resistance mechanisms, including growth rate penalties, altered metabolism, stress tolerance, and competitive ability. Resistance phenotypes that appear equivalent by AST can differ substantially in their evolutionary stability, transmissibility, and persistence, due to variation in fitness costs and compensatory evolutionary trajectories.^12–17^ Incorporating fitness assays alongside traditional AST would provide a more complete picture of bacterial behavior under selective pressure, enabling improved prediction of treatment outcomes, resistance emergence, and long-term population dynamics. In this way, fitness measurements extend susceptibility testing beyond a binary resistant vs. susceptible framework toward a more mechanistic and predictive understanding of antimicrobial resistance.

Monitoring phenotypic shifts at both individual and community levels may establish new avenues for AMR detection. Broadening the scope of measurable indicators could facilitate scalable, real-time assessments of microbial resistance. Recently, provocative new associations between antibiotic resistance and swimming motility were reported.^18–20^ Specifically, these investigators reported alterations in swimming motility in multiple Enterobacteriaceae species following acute antimicrobial exposure as well as prolonged adaptation to diverse antibiotic pressures.^18–20^ Such observations suggest a previously unrecognized mechanistic link between motility and antibiotic resistance. Swimming motility is straightforward to measure and monitor in the laboratory, and as a key fitness trait, is broadly relevant across the bacterial kingdom. Moreover, motility assays can be readily adapted for automation and miniaturization using microfluidics and microscopy. However, the implications of motility in antibiotic resistance and its potential utility as a surrogate marker for AMR remain poorly understood.

Thus, in this work, we systematically and comprehensively investigate the relationship between bacterial swimming motility and the evolution of β-lactam antibiotic resistance. Specifically, we focus on β-lactam antibiotics as they are critically important in human medicine,^21^ and have historically been used in livestock production.^22^ We employ a classic laboratory evolution approach with multiple independent replicate lineages of an *Escherichia coli (E. coli)* strain (NIST 0056) subjected to repeated β-lactam antibiotic pressure. This laboratory evolution is coupled with concurrent antimicrobial susceptibility testing, motility characterization, and whole-genome sequencing. We find evidence of a link between efflux-based or membrane-associated resistances commonly found in the environment and dysregulation of swimming motility in *E. coli,* with consistently observed mutations occurring in *marR*, *acrB*, and *frdD*/*ampC*. Further, we provide proof-of-concept that motility can serve as an early indicator of developing β-lactam resistance in both *E. coli* populations and single clones. These findings have translational potential for development of a rapid detection method for widespread AMR screening. Future work is required to determine generalizability to other bacterial strains and antibiotic classes.

## 2. Methods

### 2.1 Experimental evolution of *E. coli*

A natural isolate (avian clinical source) of *E. coli* (NIST 0056, NIST Microbial Strain Bank) was obtained from the FDA (7.1994, Penn State, Edward Dudley). The genome sequence is available in GenBank under accession CP116241.1 and is associated with BioProject PRJNA224116 and BioSample SAMN32743768. Prior to experiments, the strain underwent double colony purification to establish a clonal ancestor for five independent replicate lineages. Experimental evolution was performed by serial passage over 10 d in sterile microtiter plates as described previously.^23^ A schematic overview of this experimental workflow is provided in **Figure 1**.

**Figure 1.**
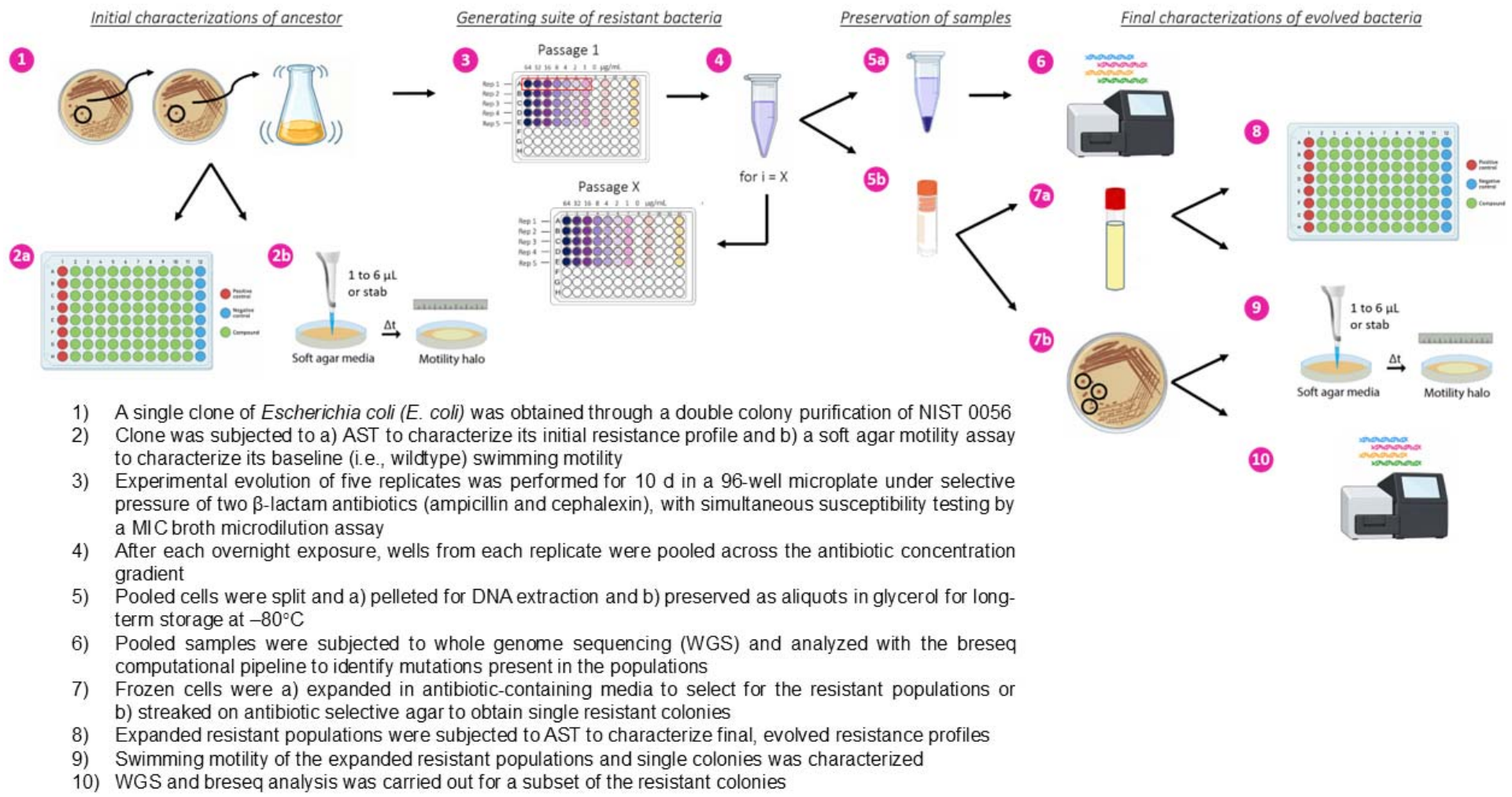
A graphical representation of the employed AMR experimental evolution workflow starting from a single clone of *Escherichia coli (E. coli)* with corresponding preservation methods and subsequent phenotypic and genotypic characterizations. Image was created using Biorender.

Briefly, the ancestral strain was cultured in a shaking incubator overnight in Luria-Bertani (LB) broth at 37 °C. Cultures were then diluted 1:1000 into 10 mL fresh LB medium, either unsupplemented or containing (0 to 512) μg/mL ampicillin or (0 to 256) μg/mL cephalexin. The diluted cultures were next transferred to 96-well plates and incubated (37 °C with orbital shaking) in a BioTek Synergy Microplate Reader (BioTek, Inc., Winooski, VT). Following each incubation period (20 h to 24 h), all wells corresponding to the full antibiotic concentration gradient for each replicate (n = 5) lineage (regardless of visible growth) were pooled. Each pooled population was diluted 1:1000 into fresh LB medium as the inoculum for the subsequent passage. Cultures were passaged daily under an increasing antibiotic concentration gradient. Aliquots of the pooled populations from each passage were archived at −80 °C (for subsequent phenotypic characterization) or pelleted and stored at −20 °C (for whole-population genome sequencing).

#### Rationale for population-based evolution

Strains were evolved as mixed populations (rather than through selection of individual clones at each passage) to maintain a large effective population size; this was done to increase the mutation supply rate and the efficiency of natural selection. Our intent was to enable competition among co-occurring variants by minimizing genetic bottlenecks and preserving standing genetic variation. For this reason, we pooled all wells across the antibiotic gradient instead of limiting selection to those at or just below the MIC. Allowing resistance to emerge through diverse mutational trajectories at the population-level provided insight into fitness dynamics and competitive interactions among resistance mechanisms. This was complemented by clonal analyses performed in parallel (*vide infra*).

### 2.2 Phenotypic reversibility

To assess phenotypic reversibility following antibiotic withdrawal, populations from selected passages (*i.e.*, corresponding to days 3, 4, 6, 7, 8, and 10) were transferred to antibiotic-free LB broth and cultured consecutively for three overnight passages. Time point selection was based on preliminary findings of varying *marR*, *acrB*, and *frdD*/*ampC* mutation frequencies at the population level. Following this antibiotic pressure withdrawal period, we subsequently performed genome sequencing and motility assays.

### 2.3 Antimicrobial susceptibility testing

Throughout the evolution experiments we monitored antibiotic resistance using minimum inhibitory concentration (MIC, defined here as the lowest antibiotic concentration to inhibit visible growth). Fold-change in the MIC was calculated relative to the ancestral MIC (day 0). Cross-resistance profiling was performed at baseline (ancestor populations) and post-selection (evolved populations) using GN4F Sensititre MIC plates (ThermoFisher). In the case of the evolved populations, frozen cells were grown to stationary phase overnight in LB supplemented with the highest antibiotic concentration at which growth was previously observed (to maintain the applied antibiotic pressure and thereby prevent any loss or reversion of the resistance phenotype). Ensuring phenotypic selection during overnight growth thus led to these assays being conducted on resistant-enriched populations rather than an entirely mixed culture. The ancestor populations were also grown to stationary phase in LB alone. As per the manufacturer’s protocol, overnight cultures were then diluted to OD = 0.1 in LB, and 30 µL of the resulting suspension was added to each well. Following this, the plates were incubated overnight (37 °C without shaking), and growth and pellet formation were assessed the following day.

### 2.4 Swimming motility assay

Frozen samples were reconstituted in selective medium (containing the highest growth-permissive antibiotic concentration for the corresponding passage) and grown overnight to stationary phase at 37 °C with orbital shaking. Cells were pelleted (8 000 × *g* for 3 min), and the supernatant decanted. One microliter of the cell pellet was used to inoculate swimming soft agar motility plates (LB containing 0.3 % Bacto agar) by stabbing the center of the plate to the center depth of the agar with a pipette tip, followed by incubation at 37 °C for 14 h, which was determined to be an optimal incubation time for resolving motility differences (**Figure S1**). Following each experiment, the diameter of the swimming zone halo was measured using the Interscience Scan 4000 automated colony counter/imager (Interscience, Saint-Nom-la-Bretèche, France), with percent reductions calculated relative to their corresponding untreated controls. Each condition was tested in triplicate. For clonal isolates (see section 2.6), three single colonies per condition were tested using the same procedure. Additional experiments were conducted in which soft agar was supplemented with antibiotics and overnight cultures were grown in non-selective media (**Figures S2-S3**).

### 2.5 Whole-population genome sequencing

Genomic DNA was extracted directly from frozen cell pellets using the NEBNext DNA extraction kit (New England Biolabs) per manufacturer’s instructions. Sequencing libraries were prepared using the NEBNext library preparation kit, followed by sequencing on an Illumina MiSeq platform using a 600-cycle V3 kit (2 × 300 bp paired-end reads). Raw sequencing reads are available in the National Center for Biotechnology Information (NCBI) Sequence Read Archive (SRA) under accession numbers SRR41192825 to SRR41192852 and are associated with BioProject PRJNA1544161. Samples were sequenced across two independent runs, comprising 17 and 11 samples, respectively. One sequencing run generated approximately 10.6 Gbp of data. The run achieved a cluster density of 731000/mm², with 96.4 % of clusters passing filter and 83.7 % of bases achieving a quality score ≥ Q30. Across both runs, raw sequencing generated approximately 22 million paired-end read pairs (≈ 6.7 Gbp), containing approximately (0.51 to 1.40) million paired-end reads (2 × 300 bp) per sample. Approximately 87 % of the reads passed quality filtering (Q ≥ 20), with average read lengths ranging from approximately (95 to 184) bp for forward reads (R1) and (130 to 193) bp for reverse reads (R2). Read quality was assessed using Phred quality scores, with the percentage of bases at Q30 or higher ranging from (91 to 96) % for R1 and 29 % for R2 in the first sequencing run, compared with (70 to 79) % for R2 in the second run. Raw reads were processed using BBduk (BBTools v38.90). Adapter sequences and low-quality bases were trimmed from both ends of reads (qtrim = rl, trimq = 20), reads containing ambiguous bases were removed (maxns = 0), and reads shorter than 100 bp after trimming were discarded (minlen = 100). Trimming statistics were recorded for each sample using BBduk log output files for quality control and reproducibility. This stringent filtering strategy ensured retention of high-confidence reads for downstream assembly and variant analysis. Following quality trimming and filtering, approximately 16 million read pairs (5 Gbp) were retained for downstream analysis. Variant calling was performed using the Breseq v.0.35.7 computational pipeline in polymorphism mode to identify mutations present in mixed populations.^24^ Junctional read evidence identified using Breseq was independently validated through splice-aware mapping with the STAR v.2.7.11b aligner.^25^ Copy number variant (CNV) detection was performed in two stages. First, coverage-based screening was performed on aligned reads by computing per-base coverage using the *depth* function in samtools v1.22.1. To reduce false positives from repetitive elements, regions annotated as transposases, insertion sequences, or repeats in the reference GFF file were excluded using the *intersect* function in bedtools v2.31.1. Next, mean coverage was calculated for each region and normalized to the median coverage across all genes in the sample. Potential duplicate regions were identified based on normalized coverage ≥ 1.8. Following screening, all potential CNVs were confirmed using the *haplocn.mops* function in the cn.MOPS v1.54.0 R package.^26^ Additional software used included PhiSpy v4.2.21^27^ to identify putative prophage sequences in the bacterial genome and AMRFinderPlus v4.2.5^28^ to identify antimicrobial resistance and virulence genes.

### 2.6 Clonal isolation and phenotypic characterization

To isolate clones from the evolved populations, samples from selected passages were streak-plated on LB agar supplemented with highest concentration of ampicillin that permitted growth during that passage. Following overnight incubation at 37 °C, three single colonies were selected from each of the five independently evolved populations and the ancestral lineages (grown without antibiotic selection). Each colony underwent: (1) swimming motility assays, performed in triplicate as described in Section 2.4, (2) preservation in an aqueous solution with a volume fraction of 25 % glycerol at −80 °C, and (3) whole-genome sequencing. For sequencing, colonies were resuspended in 200 μL PBS, then pelleted and stored at −20 °C until time of DNA extraction. Sequencing was then performed with the Illumina MiSeq platform as described above.

### 2.7 Statistical analysis

All statistical analyses were performed in R v4.5.2. To assess differences between population-level and colony-level measurements, nonparametric Wilcoxon rank-sum (Mann-Whitney U) tests were conducted separately within each replicate. This approach was chosen because population and colony measurements within replicates consisted of multiple independent observations and did not meet assumptions of normality (as assessed through visual diagnostics). For each replicate, population and colony distributions were compared independently, and two-sided tests were used in all cases. Statistical significance was assessed at α = 0.05.

## 3. Results and Discussion

### 3.1 Reversible loss of swimming motility accompanies β-lactam resistance evolution in E. coli

The ancestor clone used in this work was expanded from an environmental isolate (avian) of E. coli (NIST 0056) with an unknown history of antibiotic exposure. Upon initial antimicrobial susceptibility testing (AST), this progenitor displayed intrinsic resistance to tetracycline and the aminoglycoside drug class, but was susceptible to β-lactam antibiotics, including ampicillin and cephalexin (MIC ≤ (4 to 8) μg/mL, as outlined by the Clinical and Laboratory Standards Institute (CLSI) guidelines (**Table S1**).^29^ After 10 d of serial passaging under continuous β-lactam exposure, all five replicate populations for each antibiotic exhibited MIC increases, showing decreased susceptibility, albeit with varied magnitudes (**Figure 2a,d**). Levels of evolved resistance were categorized as follows: “Susceptible (S)” (≤ 8 μg/mL), “Intermediate (I)” (= 16 μg/mL), and “Resistant (R)” (≥ 32 μg/mL).^29^ Specifically, intermediate or resistant phenotypes emerged in four of the five replicates under exposure to each antibiotic. Notably, these lab-evolved resistant populations also exhibited cross-resistance to multiple other antibiotics (**Table S1**).

**Figure 2.**
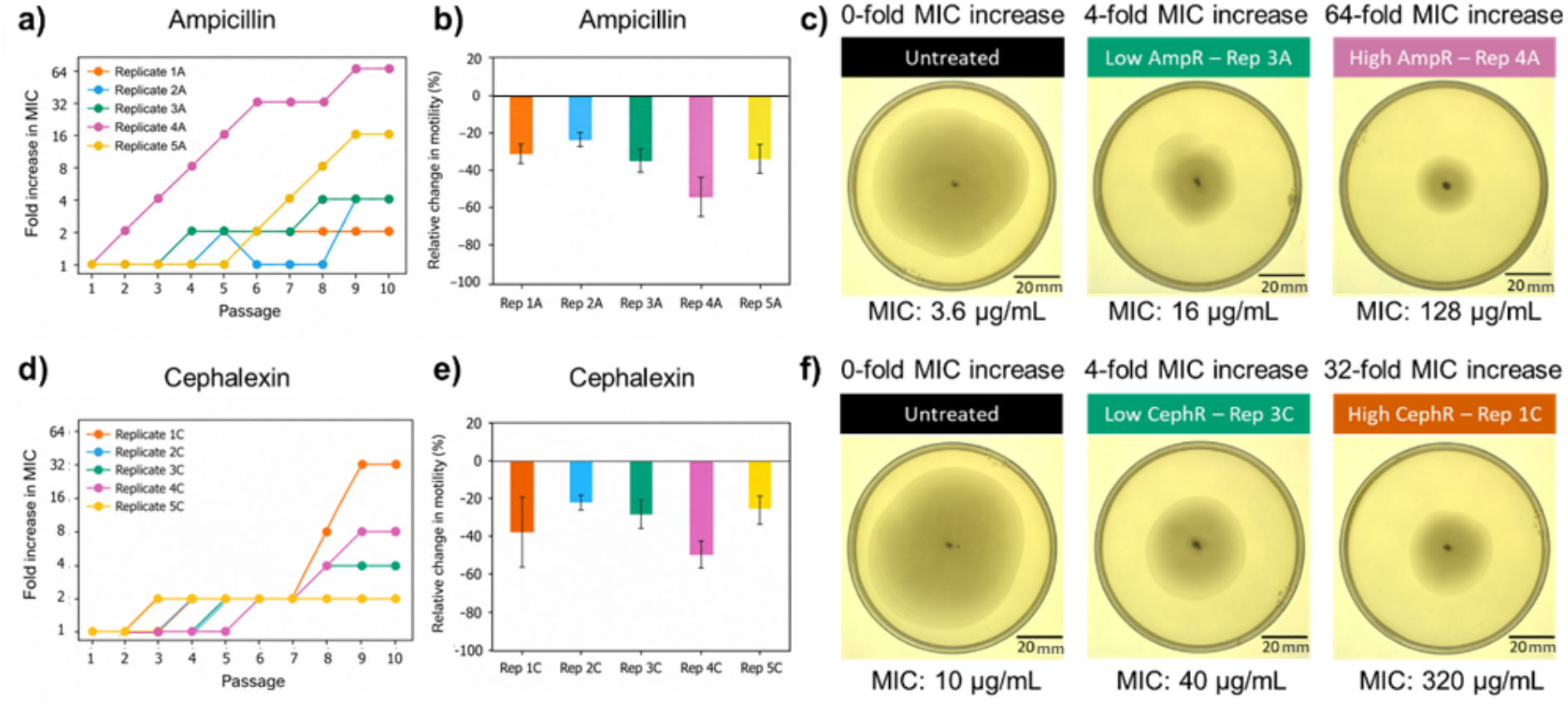
Demonstrated link between β-lactam antibiotic resistance evolution and reduction in phenotypic swimming motility in *E. coli* NIST 0056. a,d) Phenotypic resistance profiles for five independent replicate lineages of *E. coli* over 10 d of evolution under selective antibiotic pressure with increasing, subinhibitory concentrations of a) ampicillin and d) cephalexin. b,e) Swimming motility of five independent replicate lineages of *E. coli* after exposure to b) ampicillin and e) cephalexin for ten passages. c,f) Representative swim zones (raw data) in LB motility agar shown for bacterial populations passaged in LB alone for 10 d versus in the presence of c) ampicillin and f) cephalexin. In c) and f), “Low/Intermediate AmpR” and “Low/Intermediate CephR” are MIC ≤ (16 to 32) μg/mL; “High AmpR” and “High CephR” are MIC > (16 to 32) μg/mL. (*i.e.*, “Susceptible” ≤ (4 to 8) μg/mL < “Low/Intermediate R” ≤ 8 to 16 to 32 μg/mL < “High R”). We generally observe a significant reduction in (various changes in) swimming motility (as well) across all five replicates for *E. coli* exposed to both antibiotics, thereby suggesting reduced motility as a phenotypic correlate of AMR in this system. Error bars represent ± 1 standard deviation of the mean for *n = 3* independent biological replicates. ^1^ For Enterobacteriaceae, including *E. coli*, identified CLSI breakpoints for ampicillin are as follows: susceptible, ≤ (4 to 8) μg/mL; intermediate, (8 to 16) μg/mL; resistant, ≥ 32 μg/mL. CLSI breakpoints for cephalexin are not currently defined and so values were based on ampicillin.

In conjunction with evolved resistance, we observed a 20 % to 60 % reduction in bacterial swimming motility across all replicate populations exposed to ampicillin or cephalexin (**Figure 2b,e**). In these cases, the diameter of the soft agar swimming motility haloes were notably reduced in the antibiotic-exposed populations (*p* < 0.01, Kruskal-Wallis test) compared to the untreated controls (**Figure 2c,f; Table S2**). Within the ampicillin-exposed populations, the level of motility reduction differed significantly among the five replicate populations (*p* < 0.0005, Kruskal-Wallis test). Additional post hoc statistical analyses comparing replicate populations are provided in the Supplementary Information (SI) (**Tables S3-S6**). We found that greater motility losses were generally associated with higher degrees of evolved β-lactam resistance. Following removal of the antibiotic pressure, motility returned to baseline, as observed in a subset of the ampicillin-selected replicates (**Figures S4-S5, Table S7**), hinting at a transient, adaptive phenotype later confirmed by genomics (*vide infra*). This suggests that evolving and maintaining β-lactam resistance incurs fitness costs, as a reversible response to antibiotic pressure, potentially related to flagellar synthesis or function (*N.B.*: the production of flagella and resulting motility are energy-intensive processes). It is possible that motility is among the first cellular processes impacted under β-lactam stress as the cell diverts resources to the acquired resistance mechanisms, ultimately allowing it to outcompete its motile counterparts without the energy expenditure required for movement (*i.e.*, with less competition and ample resources). In the absence of antibiotic selection, fitness tradeoffs may have favored the re-expansion of motile, susceptible phenotypes and competitively displaced non-motile, resistant counterparts, while spontaneous reversion of single-base mutations to the ancestral state may also have contributed. Alternatively, motility may simply be under the same regulatory cascade as the genetic resistance mechanisms acquired (*vide infra*). Yet, there were instances where motility loss did not align with the degree of evolved resistance, such as replicate 2C where the magnitude of motility loss was less than anticipated based on the degree of resistance acquired. This observation may reflect normal biological variation among replicates, or be attributable to the presence of mixed phenotypes resulting from the resistance selection method (*i.e.*, the subsequent pooling and expansion of replicate populations across the antibiotic concentration gradient following each passage). These results nevertheless suggest a reciprocal relationship between evolved resistance and swimming motility. Similar associations between these two phenotypes across other bacterial strains and antibiotic classes have been reported.^18–20^

### 3.2 Clonal outgrowths display more polarized swimming motility phenotypes compared to their parent populations

The resistance selection method involved pooling cells exposed across the antibiotic concentration gradient prior to passaging (see step 5 in **Figure 1**), which likely selected for multiple phenotypes within each replicate population. To evaluate the consequences of this approach, we compared reductions in swimming motility between: (1) *pooled* populations, presumably containing multiple intraspecific variants (despite having been enriched for resistance in selective broth prior to the soft agar assays); and (2) *individual clones*, isolated as single resistant colonies on selective agar and expected to express more uniform resistance phenotypes. This comparison was conducted specifically for populations and clones of *E. coli* exposed to ampicillin.

Single colonies selected on ampicillin-containing agar exhibited distinct motility profiles compared to their parent pooled populations at day 10 (**Figure 3**). The clonally-derived bacteria displayed more polarized motility phenotypes overall (*p* < 0.005, Mann-Whitney): we found pronounced swimming motility losses (*i.e.*, > 70 % reduction) in replicates 3A to 5A, yet little to no change in motility (*i.e.*, < 20 % reduction) in replicates 1A and 2A. The large motility reductions in clones 3A-5A were found to be significantly different from the parent populations (*p* < 0.05, Mann-Whitney), which was not the case in clones 1A and 2A. These observations suggest motility to be a mutable trait, responsive to stressors such as antibiotic exposure. In contrast, the pooled populations displayed intermediate levels of motility reduction, ranging from 20 % to 50 %. This is consistent with the suspected phenotypic heterogeneity arising from the pooling process and reflecting the relative proportions of non-motile and motile clones within each population. The clonal isolates thus exposed a wider repertoire of swimming phenotypes linked to β lactam resistance that would otherwise be obscured.

**Figure 3.**
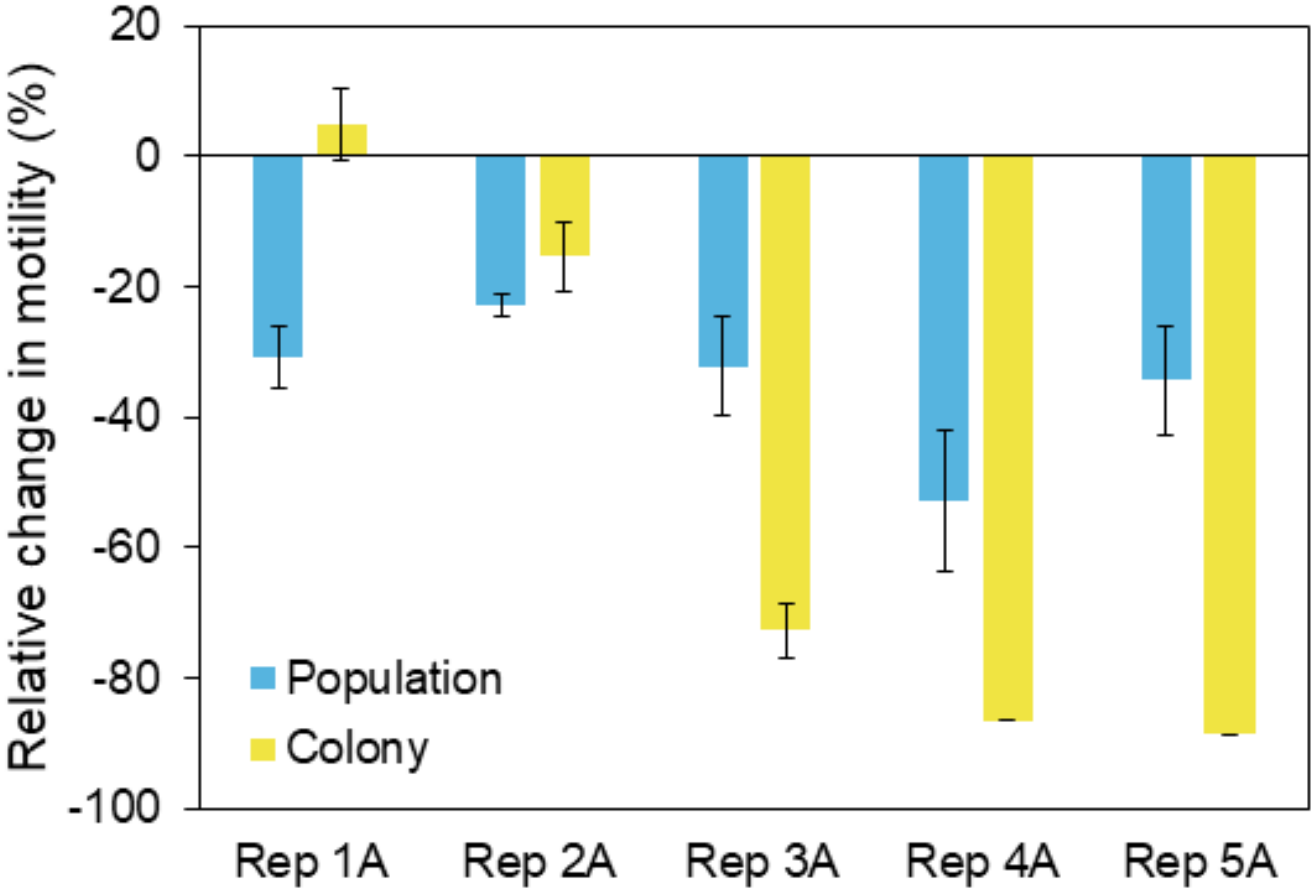
Comparison of swimming motility between replicate populations (reprinted from Figure 2b) versus single resistant colonies obtained on antibiotic selective agar (i.e., LB+ampicillin) at day 10. Specifically, three different colonies (i.e., three biological replicates) were obtained from each replicate population and run in technical triplicate (*n = 9* total for each replicate lineage) against the population-level analysis (*n = 3* for each replicate population). Thus, the average change in swimming motility of the colonies was compared to that of the population. While we observed a robust reduction in swimming motility for the replicate populations, the clonal isolates exhibited greater reductions in motility (likely associated with greater resistance), indicating that the populations may continue to harbor a non-resistant, susceptible (and more motile) phenotype/subpopulation that lowers the overall change in motility observed in the populations as compared to the clones. The reduction in swimming motility observed in the clonal isolates closely follows the level of resistance evolved in each of the replicates in Figure 2a, suggesting motility as an indicator for the *level* of resistance evolved. Error bars represent ± 1 standard deviation of the mean for both colony (*n = 9*) and population (*n = 3*) measurements.

### 3.3 Population genomics identifies mutations that converge on canonical resistance and efflux pathways under β-lactam selection

Whole-population genomics was used to identify adaptive mutations associated with the observed phenotypic trajectories. Following quality trimming, samples contained approximately 0.36 to 1.01 million paired-end read pairs (2 × 300 bp), with minimum read lengths of 100 bp (average read lengths of (141 to 201) bp for forward reads and (136 to 195) bp for reverse reads). Across the biological samples, (99 to 100) % of bases had quality scores ≥ Q20, and (94 to 99) % had quality scores ≥ Q30. After filtering, the resulting datasets provided approximately 30× to 60× genome coverage for downstream variant analysis.

Genomics analysis confirmed that the ancestor strain harbored a portfolio of chromosomal and plasmid-borne resistance determinants (**Table S8**). As anticipated from the baseline AST profile, this isolate possessed copies of *tetA*/*tetR* (encoding a tetracycline efflux pump and its repressor), along with multiple aminoglycoside-modifying genes (*aphA*, *yokD,* and *aac(3)-VIa*). The ancestor also carried the chromosomal class C β-lactamase (*ampC*),^30^ yet remained phenotypically susceptible to the β-lactams introduced. The *ampC* gene encodes an Ambler class C cephalosporinase that hydrolyzes many penicillins and early generation cephalosporins and is not inhibited by sulbactam (the β-lactamase inhibitor paired with ampicillin in the AST assay).^31,32^ This locus is typically regulated via the AmpR/AmpC axis;^33^ however, the NIST 0056 strain lacked *ampR*, suggesting limited *ampC* expression under basal conditions. These findings are consonant with those of Nong *et al.* who reported that *E. coli* K-12 MG1655, like NIST 0056, lacks *ampR* and exhibits constrained *ampC* expression in the absence of β lactam selection.^34^

In the evolved strains, sequencing revealed mutations affecting several paradigmatic AMR pathways (**Table 1**, **Figures 4-5**). Detailed genomics analysis identified a recurring deletion (Δ127 bp) in *marR* that reached fixation by day 10 in independent isolates 2C, 4A, and 5A, and occurred at high frequency (> 75 %) in replicate populations 1C, 3C, and 4C. A distinct *marR* deletion (Δ56 bp) was detected in 58 % of the replicate 3A population at day 10. MarR functions as a transrepressor positioned upstream of the multiple antibiotic resistance gene *marA*, a central regulator governing the expression of a large AMR gene network,^35^ including the AcrAB-TolC multidrug efflux system.^36,37^ Thus, large *marR* deletions may derepress AcrAB activity and potentiate antibiotic efflux, reducing intracellular antibiotic concentrations.^36–38^ Single nucleotide polymorphisms (SNPs) were also detected in *acrB* (a subunit of the AcrAB-TolC efflux complex) in eight of the ten replicates, reaching fixation in all but one (replicate 4C), with no *acrB* variants being observed in replicates 2A or 5C. Of note, 70 % (7 of 10) of lineages converged on *acrB* mutations within the PN1 subdomain of the encoded protein, reproducing the same Y49S and V127G variants shown independently to modulate drug binding energetics and efflux.^39^ Specifically, these PN1 mutations would enhance efflux by remodeling entrance to the transporter’s proximal binding pocket, facilitating substrate capture and passage through the AcrB export cycle.

**Figure 4.**
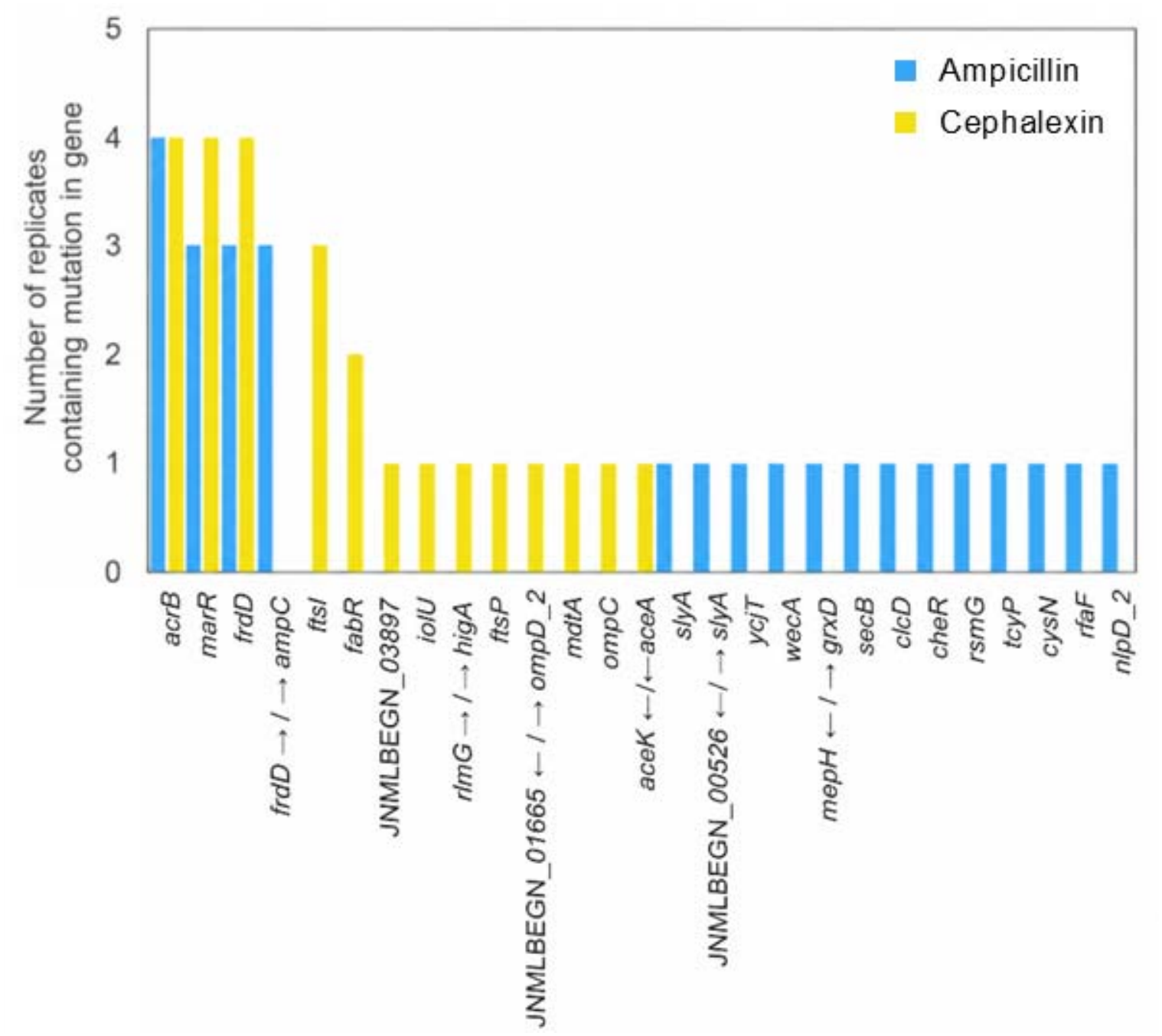
Frequencies of mutations and indels observed across genes in five replicates exposed to ampicillin and cephalexin for 10 d. **Note:** Arrows indicate the direction of the mutation or indel. If a single gene name is shown, the mutation/indel occurs within that gene. If two gene names are shown separated by a solidus, the mutation/indel lies in the intergenic region between them, with arrows indicating its direction relative to the adjacent genes.

**Figure 5.**
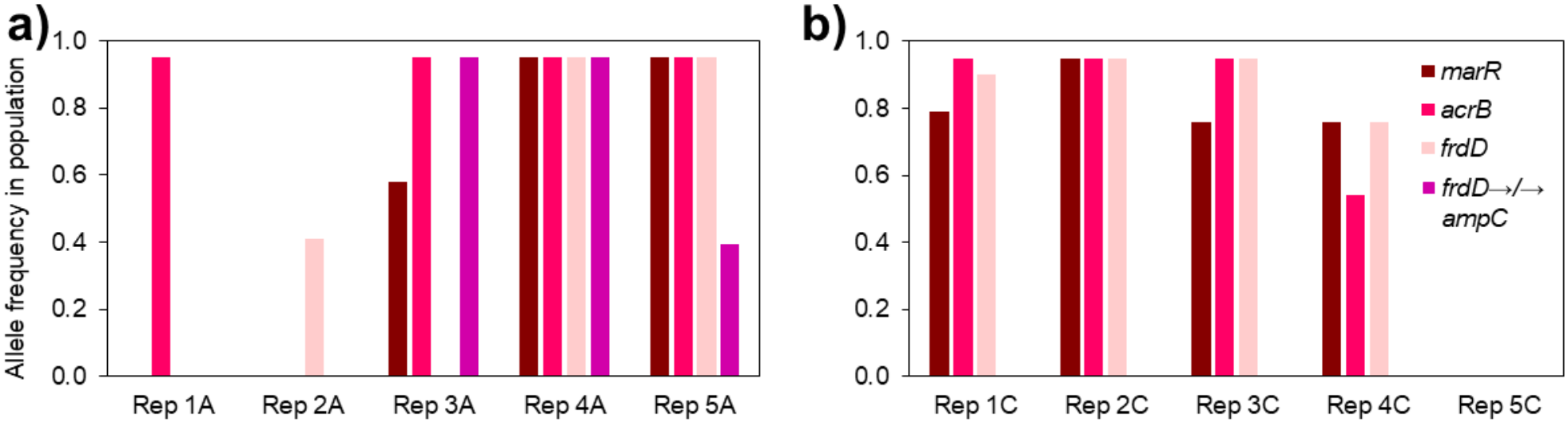
Frequencies of the four major identified mutations that swept to fixation or near fixation in replicate mixed populations after 10 d of repeated exposure to (a) ampicillin and (b) cephalexin.

**Table 1.**
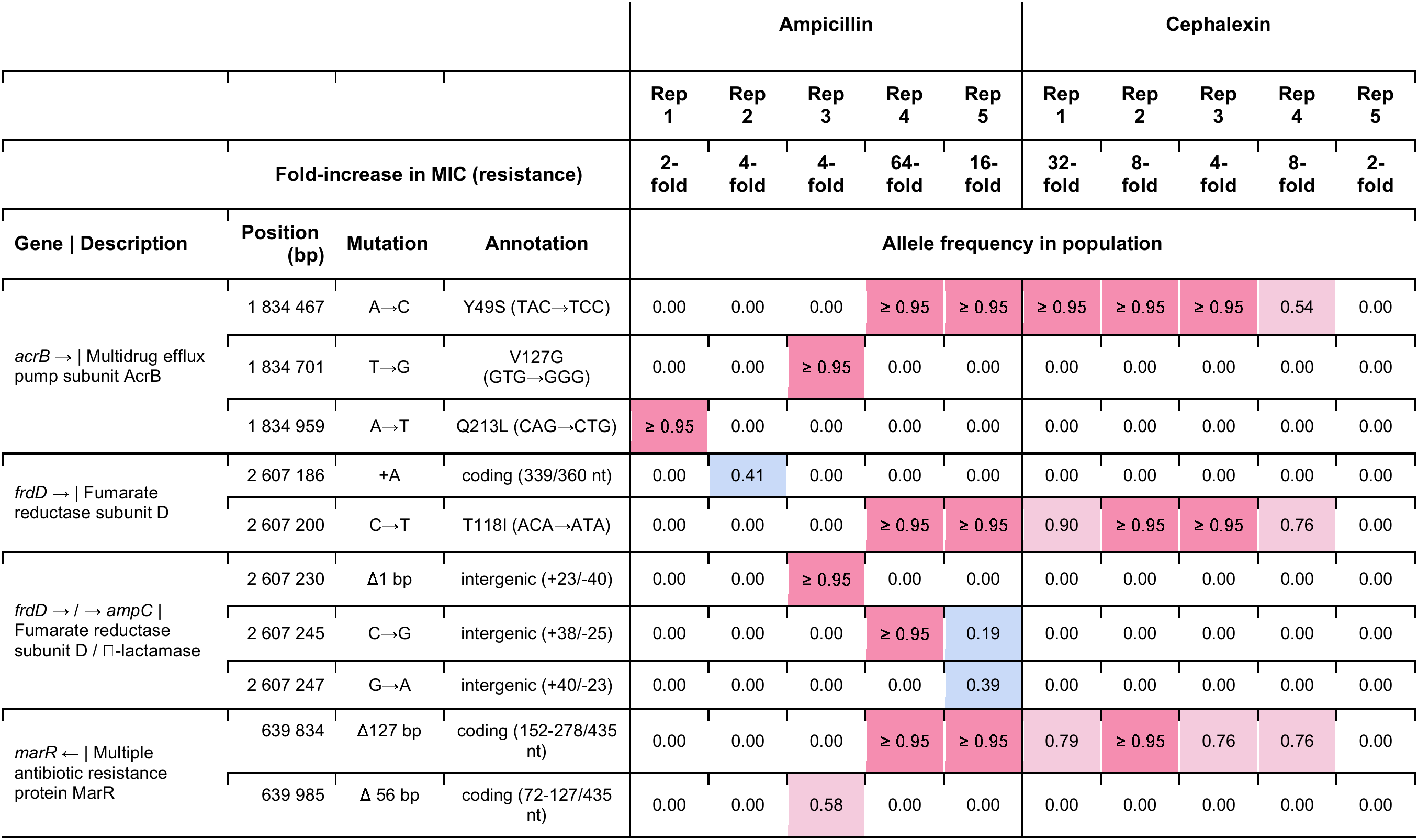
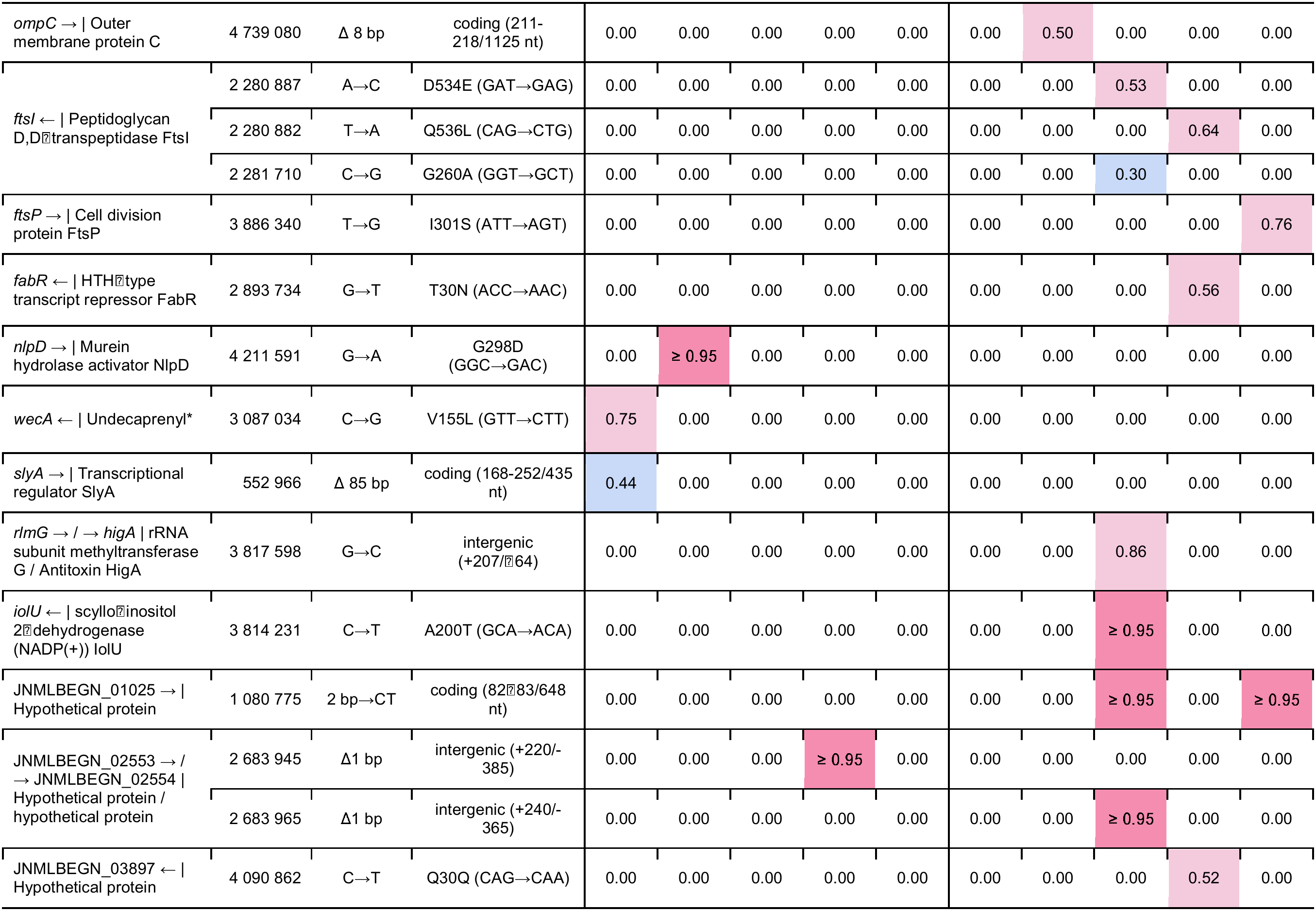
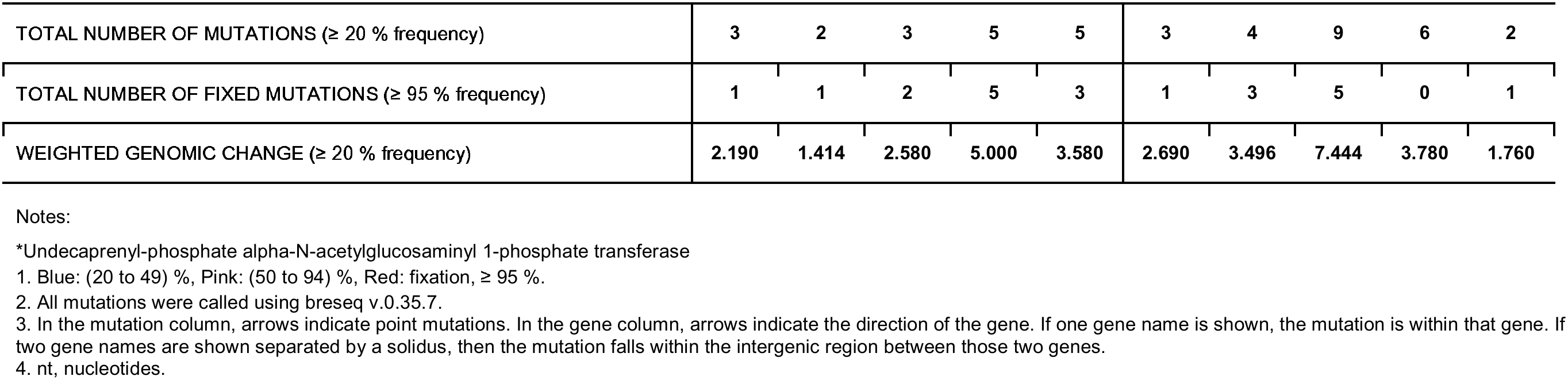
Observed mutations across replicates in avian *E. coli* after 10 passages in ampicillin and cephalexin. Mutations occurring at > 40 % frequency in at least one population and not present in the untreated control are included. Certain mutations (and their frequencies in the population) correlate with the largest increases in MIC, not with the total number of mutations. Similar mutations appear across replicates, notably in the multidrug efflux pump subunit AcrB, fumarate reductase subunit D/β-lactamase, and the multiple antibiotic resistance protein MarR.

| **Gene \| Description** | **Position (bp)** | **Mutation** | **Annotation** | **Ampicillin** |  |  |  |  | **Cephalexin** |  |  |  |  |
|  |  |  |  | **Rep 1** | **Rep 2** | **Rep 3** | **Rep 4** | **Rep 5** | **Rep 1** | **Rep 2** | **Rep 3** | **Rep 4** | **Rep 5** |
|  | **Fold-increase in MIC (resistance)** |  |  | **2-fold** | **4-fold** | **4-fold** | **64-fold** | **16-fold** | **32-fold** | **8-fold** | **4-fold** | **8-fold** | **2-fold** |
|  |  |  |  | **Allele frequency in population** |  |  |  |  |  |  |  |  |  |
|---|---|---|---|---|---|---|---|---|---|---|---|---|---|
| *acrB* → \| Multidrug efflux pump subunit AcrB | 1 834 467 | A→C | Y49S (TAC→TCC) | 0.00 | 0.00 | 0.00 | ≥ 0.95 | ≥ 0.95 | ≥ 0.95 | ≥ 0.95 | ≥ 0.95 | 0.54 | 0.00 |
|  | 1 834 701 | T→G | V127G (GTG→GGG) | 0.00 | 0.00 | ≥ 0.95 | 0.00 | 0.00 | 0.00 | 0.00 | 0.00 | 0.00 | 0.00 |
|  | 1 834 959 | A→T | Q213L (CAG→CTG) | ≥ 0.95 | 0.00 | 0.00 | 0.00 | 0.00 | 0.00 | 0.00 | 0.00 | 0.00 | 0.00 |
| *frdD* → \| Fumarate reductase subunit D | 2 607 186 | +A | coding (339/360 nt) | 0.00 | 0.41 | 0.00 | 0.00 | 0.00 | 0.00 | 0.00 | 0.00 | 0.00 | 0.00 |
|  | 2 607 200 | C→T | T118I (ACA→ATA) | 0.00 | 0.00 | 0.00 | ≥ 0.95 | ≥ 0.95 | 0.90 | ≥ 0.95 | ≥ 0.95 | 0.76 | 0.00 |
| *frdD* → / → *ampC* \| Fumarate reductase subunit D / □-lactamase | 2 607 230 | Δ1 bp | intergenic (+23/-40) | 0.00 | 0.00 | ≥ 0.95 | 0.00 | 0.00 | 0.00 | 0.00 | 0.00 | 0.00 | 0.00 |
|  | 2 607 245 | C→G | intergenic (+38/-25) | 0.00 | 0.00 | 0.00 | ≥ 0.95 | 0.19 | 0.00 | 0.00 | 0.00 | 0.00 | 0.00 |
|  | 2 607 247 | G→A | intergenic (+40/-23) | 0.00 | 0.00 | 0.00 | 0.00 | 0.39 | 0.00 | 0.00 | 0.00 | 0.00 | 0.00 |
| *marR* ← \| Multiple antibiotic resistance protein MarR | 639 834 | Δ127 bp | coding (152-278/435 nt) | 0.00 | 0.00 | 0.00 | ≥ 0.95 | ≥ 0.95 | 0.79 | ≥ 0.95 | 0.76 | 0.76 | 0.00 |
|  | 639 985 | Δ 56 bp | coding (72-127/435 nt) | 0.00 | 0.00 | 0.58 | 0.00 | 0.00 | 0.00 | 0.00 | 0.00 | 0.00 | 0.00 |
| *ompC* → \| Outer membrane protein C | 4 739 080 | Δ 8 bp | coding (211-218/1125 nt) | 0.00 | 0.00 | 0.00 | 0.00 | 0.00 | 0.00 | 0.50 | 0.00 | 0.00 | 0.00 |
| *ftsI* ← \| Peptidoglycan D,D□transpeptidase FtsI | 2 280 887 | A→C | D534E (GAT→GAG) | 0.00 | 0.00 | 0.00 | 0.00 | 0.00 | 0.00 | 0.00 | 0.53 | 0.00 | 0.00 |
|  | 2 280 882 | T→A | Q536L (CAG→CTG) | 0.00 | 0.00 | 0.00 | 0.00 | 0.00 | 0.00 | 0.00 | 0.00 | 0.64 | 0.00 |
|  | 2 281 710 | C→G | G260A (GGT→GCT) | 0.00 | 0.00 | 0.00 | 0.00 | 0.00 | 0.00 | 0.00 | 0.30 | 0.00 | 0.00 |
| *ftsP* → \| Cell division protein FtsP | 3 886 340 | T→G | I301S (ATT→AGT) | 0.00 | 0.00 | 0.00 | 0.00 | 0.00 | 0.00 | 0.00 | 0.00 | 0.00 | 0.76 |
| *fabR* ← \| HTH□type transcript repressor FabR | 2 893 734 | G→T | T30N (ACC→AAC) | 0.00 | 0.00 | 0.00 | 0.00 | 0.00 | 0.00 | 0.00 | 0.00 | 0.56 | 0.00 |
| *nlpD* → \| Murein hydrolase activator NlpD | 4 211 591 | G→A | G298D (GGC→GAC) | 0.00 | ≥ 0.95 | 0.00 | 0.00 | 0.00 | 0.00 | 0.00 | 0.00 | 0.00 | 0.00 |
| *wecA* ← \| Undecaprenyl\* | 3 087 034 | C→G | V155L (GTT→CTT) | 0.75 | 0.00 | 0.00 | 0.00 | 0.00 | 0.00 | 0.00 | 0.00 | 0.00 | 0.00 |
| *slyA* → \| Transcriptional regulator SlyA | 552 966 | Δ 85 bp | coding (168-252/435 nt) | 0.44 | 0.00 | 0.00 | 0.00 | 0.00 | 0.00 | 0.00 | 0.00 | 0.00 | 0.00 |
| *rlmG* → / → *higA* \| rRNA subunit methyltransferase G / Antitoxin HigA | 3 817 598 | G→C | intergenic (+207/□64) | 0.00 | 0.00 | 0.00 | 0.00 | 0.00 | 0.00 | 0.00 | 0.86 | 0.00 | 0.00 |
| *iolU* ← \| scyllo□inositol 2□dehydrogenase (NADP(+)) IolU | 3 814 231 | C→T | A200T (GCA→ACA) | 0.00 | 0.00 | 0.00 | 0.00 | 0.00 | 0.00 | 0.00 | ≥ 0.95 | 0.00 | 0.00 |
| JNMLBEGN_01025 → \| Hypothetical protein | 1 080 775 | 2 bp→CT | coding (82□83/648 nt) | 0.00 | 0.00 | 0.00 | 0.00 | 0.00 | 0.00 | 0.00 | ≥ 0.95 | 0.00 | ≥ 0.95 |
| JNMLBEGN_02553 → / → JNMLBEGN_02554 \| Hypothetical protein / hypothetical protein | 2 683 945 | Δ1 bp | intergenic (+220/-385) | 0.00 | 0.00 | 0.00 | ≥ 0.95 | 0.00 | 0.00 | 0.00 | 0.00 | 0.00 | 0.00 |
|  | 2 683 965 | Δ1 bp | intergenic (+240/-365) | 0.00 | 0.00 | 0.00 | 0.00 | 0.00 | 0.00 | 0.00 | ≥ 0.95 | 0.00 | 0.00 |
| JNMLBEGN_03897 ← \| Hypothetical protein | 4 090 862 | C→T | Q30Q (CAG→CAA) | 0.00 | 0.00 | 0.00 | 0.00 | 0.00 | 0.00 | 0.00 | 0.00 | 0.52 | 0.00 |
| TOTAL NUMBER OF MUTATIONS (≥ 20 % frequency) |  |  |  | **3** | **2** | **3** | **5** | **5** | **3** | **4** | **9** | **6** | **2** |
| TOTAL NUMBER OF FIXED MUTATIONS (≥ 95 % frequency) |  |  |  | **1** | **1** | **2** | **5** | **3** | **1** | **3** | **5** | **0** | **1** |
| WEIGHTED GENOMIC CHANGE (≥ 20 % frequency) |  |  |  | **2.190** | **1.414** | **2.580** | **5.000** | **3.580** | **2.690** | **3.496** | **7.444** | **3.780** | **1.760** |

Notes:
\*Undecaprenyl-phosphate alpha-N-acetylglucosaminyl 1-phosphate transferase
1. Blue: (20 to 49) %, Pink: (50 to 94) %, Red: fixation, ≥ 95 %.
2. All mutations were called using breseq v.0.35.7.
3. In the mutation column, arrows indicate point mutations. In the gene column, arrows indicate the direction of the gene. If one gene name is shown, the mutation is within that gene. If two gene names are shown separated by a solidus, then the mutation falls within the intergenic region between those two genes.
4. nt, nucleotides.

Multiple mutations were also detected in the fumarate reductase subunit D (*frdD),* a membrane-anchoring protein within the fumarate reductase (FrdABCD) complex, including a nonsynonymous substitution (T118I) that reached fixation in replicates 4A, 5A, 2C, and 3C, and exceeded 75 % prevalence in replicates 1C and 4C. These mutations likely reflect metabolic and membrane-associated, compensatory adaptations that mitigate β-lactam-induced envelope stress and reduce growth-dependent killing. These findings are consistent with prior reports linking *frdD* mutations to amoxicillin adaptation in *E. coli*.^40^ Interestingly, mutations in the intergenic region between *frdD* and *ampC* (encompassing the *ampC* promoter) were exclusively observed among ampicillin-selected populations, with frequencies ranging from 18.5 % to reaching fixation, in three of five replicates. These latter results resembled the early-phase resistance mechanism described by Nong et al., where stepwise amoxicillin exposure in *E. coli* MG1655 fostered the accrual of transcription-enhancing *ampC* promoter mutations, presaging later gene amplifications.^34^ The selective *ampC* transcriptional remodeling under penicillin pressure likely reflects a stronger fitness benefit than under cephalexin exposure. AmpC efficiently hydrolyzes ampicillin^31^ and cephalexin,^41^ despite the latter’s greater intrinsic ring stability relative to many classical cephalosporins.^42^ This suggests that catalytic capacity alone is unlikely to explain the diverging responses; rather, the limited *ampC* remodeling under cephalexin indicates diversion of adaptation strategies toward alternative routes. Accordingly, only the cephalexin-exposed populations exhibited point mutations in the transpeptidase domain of FtsI (a β-lactam target that catalyzes cell wall peptidoglycan cross linking), including D534E (53 % in replicate 3C) and Q536L (64 % in replicate 4C); in the cell division protein FtsP, including I301S (76 % in replicate 5C); and in the fatty acid biosynthesis repressor FabR (playing a role in lipid membrane permeability), including T30N (56 % in replicate 4C). These substitutions are likely to diminish β-lactam binding and/or alter drug entry, implicating target-site remodeling as a possible evolutionary trajectory in response to cephalexin.

While the detected sequence variants provided insight into several adaptive pathways, resistance can also arise through gene amplification. We therefore also performed copy number variation (CNV) analysis to identify additional events that may have contributed to the observed resistant phenotypes. CNV analysis of the day 10 samples showed increased read coverage consistent with segmental amplification of a large 149.5 kbp region (≈ 140 coding sequences), including the *acrAB* efflux operon, in replicates 1A and (to a lesser extent) 3A (**Figure S6**). A second amplified segment of 243.5 kbp was observed in all five ampicillin-exposed replicates as well as replicate 4C, comprising roughly 220 coding sequences and including *ampC* (**Figure S6**). In parallel, we also detected focal copy number attenuation involving a 52.8 kbp segment (spanning 42 predicted genes) in replicates 2A and 1C at day 10. This latter interval bore phage-like features and was delimited by att-like 14 bp direct repeats (GCAGAACACATTAA), possibly representing a lambdoid prophage locus excised under antibiotic stress, likely as a way to reduce metabolic burden and improve fitness (**Figure S6**).

The presence of these amplifications (particularly the 243.5 kbp region) across multiple independent replicates under ampicillin pressure suggests a non-random process. Interestingly, both amplified regions were bracketed by copies of the insertion sequence (IS) 110 family transposase, *IS621*.^43^ The recurrence of *IS621* at the boundaries of both regions raises the possibility that mobile genetic elements contributed to the amplifications, potentially through homologous recombination between IS copies or transposition-like events.^43,44^ While the precise mechanisms await clarification, these findings accord with a more general role for IS elements in facilitating large-scale genomic amplifications under selective pressure.^45^

Taken together, the efflux-associated *acrB* mutations align with the large deletions detected in *marR*, supporting a multi-tiered strategy to increase efflux capacity. More broadly, these patterns suggest that cells may initially develop resistance by increasing efflux through rapid and ubiquitous—though energetically costly—means of reducing intracellular antibiotic exposure. Such strategies are inherently pleiotropic and epistatic, affecting not only resistance but also cellular energetics and physiology. In this context, mutations in *frdD* appear epistatic with *acrB*, *marR*, and *ampC*, potentially helping to buffer fitness costs associated with elevated efflux and cell envelope stress. Consistent with this, reduced motility associated with *frdD* and efflux regulation may represent a collateral consequence of selection on efflux and regulatory pathways rather than a directly selected trait. Motility specifically appears to track the presence of a *marR* deletion; in replicates 1A, 2A, and 5C, no motility reduction was observed, and no *marR* deletion occurred. An *acrB* mutation alone (i.e., a local change) does not appear sufficient to impact motility or induce pleiotropic effects, suggesting that *marR* deletion (global regulation) may be required alongside *acrB*, hinting again at a potential epistatic relationship.

In parallel, we observed amplification of *ampC*, indicating increased locus abundance as an additional mechanism and layered strategy contributing to β-lactam resistance. Unlike point mutations, which alter protein function, gene amplifications increase dosage, and bacteria frequently deploy both strategies simultaneously. In this case, change-of-function appears to confer resistance more rapidly (replicate 4A, reaching 64-fold increases in MIC, *R-Amp*) compared to increased gene dosage through genetic amplification (replicate 3A, reaching 4-fold increases in MIC, *I-Amp*). The co-occurrence of *ampC* amplification with metabolic mutations (*e.g.*, *frdD*) lends further evidence for epistatic coupling, with β-lactamase activity likely supported by the same compensatory adaptations described above. Together, these results indicate that β-lactam exposure selects for coordinated changes in metabolism, membrane energetics, and efflux capacity, collaboratively shaping resistance and tolerance phenotypes.

These results also raise the possibility for regulatory cross-talk between efflux and motility systems, such as that documented previously in *Salmonella enterica*.^46^ In that species, MarA (with SoxS, Rob, and RamA) attenuated flagellar gene expression (and hence motility) through a master regulator, FlhDC. Conservation of the *mar*-*sox*-*rob* regulon among Enterobacteriaceae, along with the high energy cost of flagellar synthesis, suggests a potential regulatory link between *marA* derepression, efflux remodeling, and impaired motility. A parallel mechanism has been reported in *E. coli*, where AcrR (a repressor of the *acrAB* operon) may also repress motility through the *flhDC* promoter.^47^ Although no *acrR* mutations were detected in our evolved strains, altered AcrR expression or activity cannot be excluded as a contributor to the observed motility changes.

### 3.4 Longitudinal genomic analysis reveals distinct evolutionary trajectories of resistance that point towards a conserved motility response

To investigate the longitudinal trends in antibiotic resistance relative to motility, we next characterized swimming behavior in two replicate lineages—replicates 3A and 4A, corresponding to Intermediate (I-Amp; MIC = 16 μg/mL) and Resistant (R-Amp; MIC = 128 μg/mL) phenotypes, respectively (**Figure 2a**)—across multiple time points along their distinct evolutionary trajectories. Replicate 3A exhibited modest MIC increases over time (*i.e.*, 2-fold to 4-fold from passages 4 to 10), yet showed significant reductions in motility, particularly at later passages (7 to 10) in the clonal populations relative to the mixed populations (**Figure 6b**). In contrast, replicate 4A showed substantial early MIC increases (*i.e.*, 2-fold to 64-fold from passages 2 to 10, including four successive stepwise increments), but exhibited less pronounced reductions in motility at earlier time points than would be expected if motility scaled with MIC (**Figure 6a**), with clonal outgrowths mirroring population-level motility changes until day 10.

**Figure 6.**
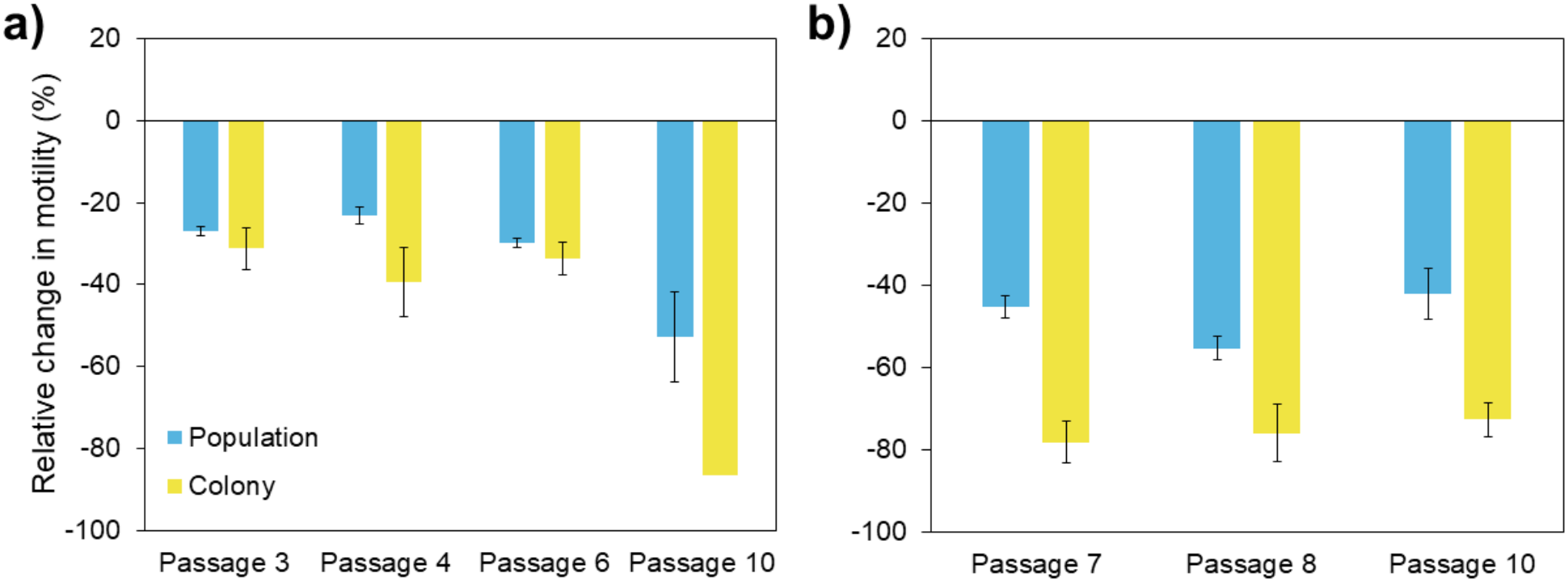
Characterization of swimming motility dynamics over successive passage numbers. Swimming motility of a) high AmpR (replicate 4A, purple points in Fig. 2a) and b) low AmpR (replicate 3A, green points in Fig. 2a) *E. coli* strains at different passages along their evolutionary paths. Passages were selected based on expression of various resistant phenotypes occurring at those time points (as indicated by different fold-increases in the MIC shown in Figure 2a). Though the absolute magnitude of reduction in motility is different over time and between replicates, there was concordance in motility reduction for all resistant phenotypes. Interestingly, however, the reduction in motility does not seem to correspond to passage number or to the resulting level of resistance at that passage for either replicate (i.e., reduction in motility does not appear to change over time, even though MIC/resistance levels are changing over time). This observation could thus suggest using motility as an early indicator or proxy for AMR, with the ability to detect phenotypic changes occurring in the population at or before the onset of AMR. Error bars represent ± 1 standard deviation of the mean for both colony (*n = 9*) and population (*n = 3*) measurements.

Time-course genomics analysis confirmed that the evolutionary paths of the two lineages were distinct (**Figure 7**). In replicate 3A, canonical resistance-associated mutations in *acrB, frdD, ampC,* and *marR* were not detected prior to day 7 (**Table 2**). Instead, mutations in non-canonical AMR-associated genes, (*pglA, ycjT*, and an uncharacterized protein) rose to fixation by day 3 and were maintained at 100 % frequency until days 7 to 8, after which they were completely lost from the population (**Figure S7**). Although none of these genes are classical resistance determinants, they are likely linked to cell envelope structure, membrane properties, metabolism, stress tolerance, and/or gene regulation—traits that influence fitness under antibiotic stress without conferring high-level resistance. By days 8 to 10, however, mutations in *acrB, frdD/ampC,* and *marR* were detected at 100 % frequency, converging toward a more conventional resistance path. These genomic trends align with the slow development of resistance observed in replicate 3A (**Figure 2a**). In contrast, replicate 4A exhibited rapid emergence of mutations in *acrB, frdD,* and *marR,* which reached substantial frequencies in the population by day 4 (**Table 3**), and expanded to fixation by day 6. These genomic dynamics were consistent with the rapid, stepwise increase in phenotypic resistance observed in replicate 4A (**Figure 2a**).

**Figure 7.**
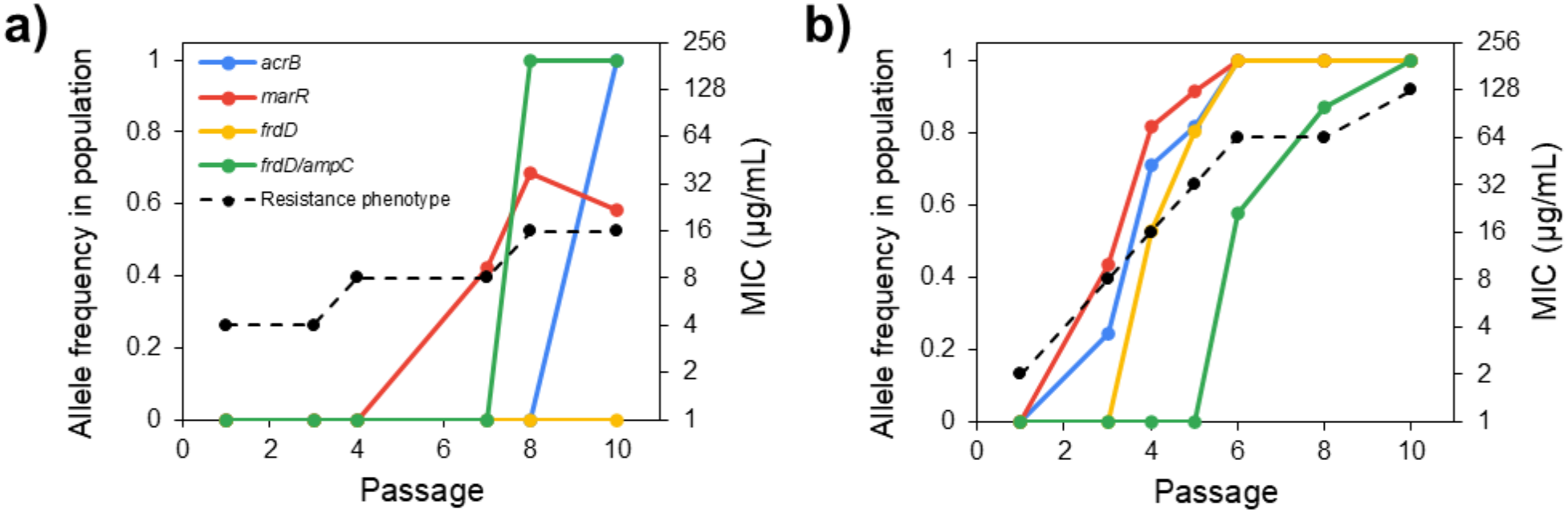
Two distinct evolutionary trajectories in individual replicates and their associated effects on phenotypic resistance, as measured by changes in MIC: a) I-Amp (Replicate 3A) and b) R-Amp (Replicate 4A). Divergent evolutionary trajectories in two individual replicates lead to distinct resistance phenotypes, as reflected by changes in MIC.

**Table 2.**
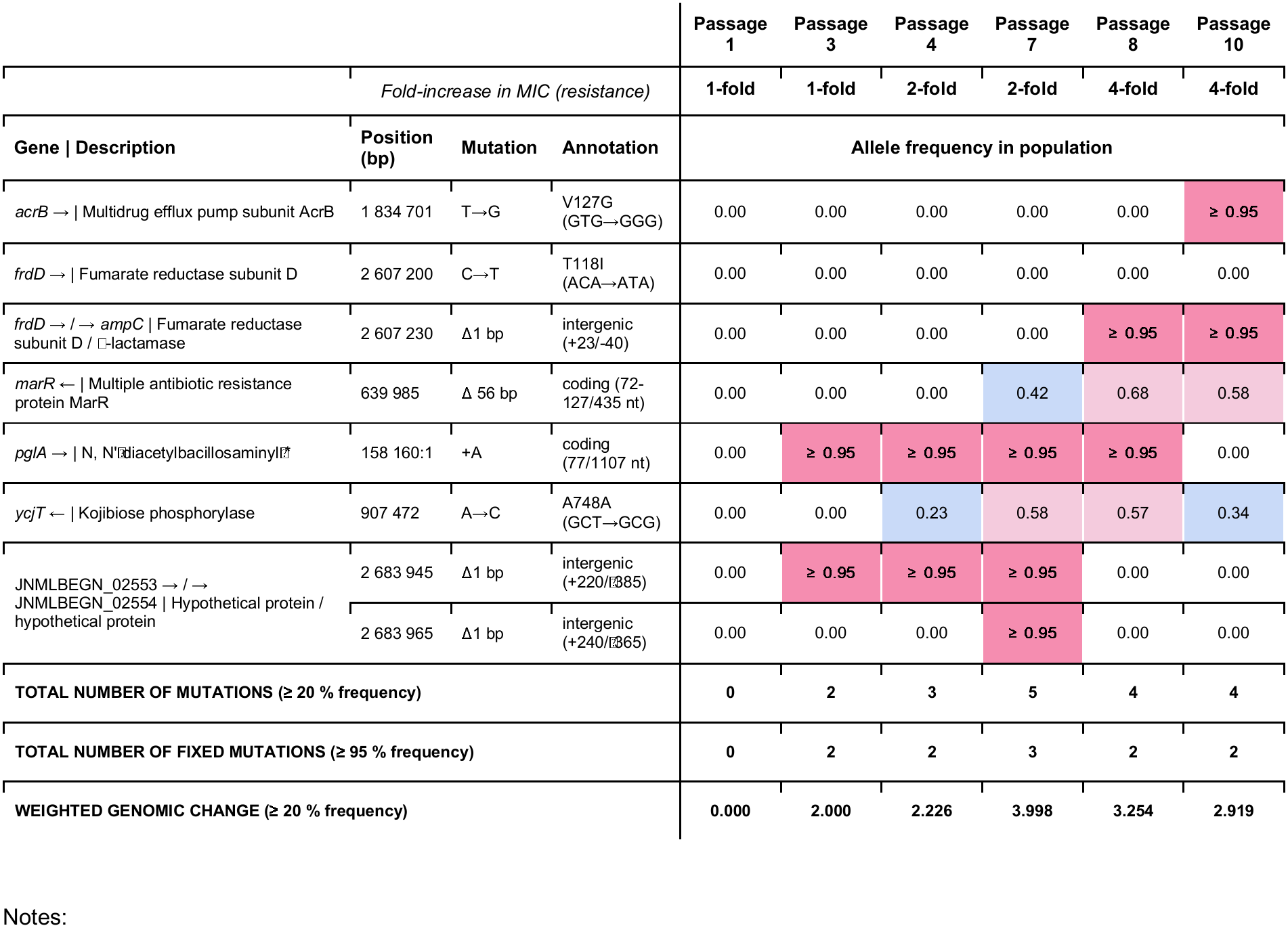
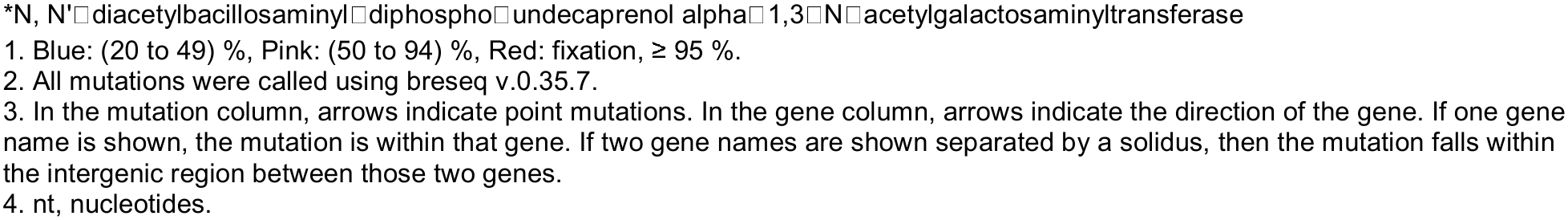
Observed mutations across passage in low I-Amp avian E. coli (replicate 3, Figure 1). Mutations appearing at > 40 % frequency in at least one of the populations and not present in the untreated control are included. It appears certain mutations fix in the population initially and then are lost in order for other mutations (e.g., *acrB*, fumarate reductase subunit D/β-lactamase) to arise and fix in order to gain resistance. If the wrong mutations are picked up initially, as it appears to be the case in this lineage, resistance is slower to emerge.

|  |  |  |  | **Passage 1** | **Passage 3** | **Passage 4** | **Passage 7** | **Passage 8** | **Passage 10** |
|  | *Fold-increase in MIC (resistance)* |  |  | **1-fold** | **1-fold** | **2-fold** | **2-fold** | **4-fold** | **4-fold** |
| **Gene \| Description** | **Position (bp)** | **Mutation** | **Annotation** | **Allele frequency in population** |  |  |  |  |  |
|---|---|---|---|---|---|---|---|---|---|
| *acrB* → \| Multidrug efflux pump subunit AcrB | 1 834 701 | T→G | V127G (GTG→GGG) | 0.00 | 0.00 | 0.00 | 0.00 | 0.00 | ≥ 0.95 |
| *frdD* → \| Fumarate reductase subunit D | 2 607 200 | C→T | T118I (ACA→ATA) | 0.00 | 0.00 | 0.00 | 0.00 | 0.00 | 0.00 |
| *frdD* → / → *ampC* \| Fumarate reductase subunit D / □-lactamase | 2 607 230 | Δ1 bp | intergenic (+23/-40) | 0.00 | 0.00 | 0.00 | 0.00 | ≥ 0.95 | ≥ 0.95 |
| *marR* ← \| Multiple antibiotic resistance protein MarR | 639 985 | Δ 56 bp | coding (72-127/435 nt) | 0.00 | 0.00 | 0.00 | 0.42 | 0.68 | 0.58 |
| *pglA* → \| N, N'□diacetylbacillosaminyl□\* | 158 160:1 | +A | coding (77/1107 nt) | 0.00 | ≥ 0.95 | ≥ 0.95 | ≥ 0.95 | ≥ 0.95 | 0.00 |
| *ycjT* ← \| Kojibiose phosphorylase | 907 472 | A→C | A748A (GCT→GCG) | 0.00 | 0.00 | 0.23 | 0.58 | 0.57 | 0.34 |
| JNMLBEGN_02553 → / → JNMLBEGN_02554 \| Hypothetical protein / hypothetical protein | 2 683 945 | Δ1 bp | intergenic (+220/□385) | 0.00 | ≥ 0.95 | ≥ 0.95 | ≥ 0.95 | 0.00 | 0.00 |
|  | 2 683 965 | Δ1 bp | intergenic (+240/□365) | 0.00 | 0.00 | 0.00 | ≥ 0.95 | 0.00 | 0.00 |
| **TOTAL NUMBER OF MUTATIONS (≥ 20 % frequency)** |  |  |  | **0** | **2** | **3** | **5** | **4** | **4** |
| **TOTAL NUMBER OF FIXED MUTATIONS (≥ 95 % frequency)** |  |  |  | **0** | **2** | **2** | **3** | **2** | **2** |
| **WEIGHTED GENOMIC CHANGE (≥ 20 % frequency)** |  |  |  | **0.000** | **2.000** | **2.226** | **3.998** | **3.254** | **2.919** |

Notes:
\*N, N'□diacetylbacillosaminyl□diphospho□undecaprenol alpha□1,3□N□acetylgalactosaminyltransferase
1. Blue: (20 to 49) %, Pink: (50 to 94) %, Red: fixation, ≥ 95 %.
2. All mutations were called using breseq v.0.35.7.
3. In the mutation column, arrows indicate point mutations. In the gene column, arrows indicate the direction of the gene. If one gene name is shown, the mutation is within that gene. If two gene names are shown separated by a solidus, then the mutation falls within the intergenic region between those two genes.
4. nt, nucleotides.

**Table 3.**
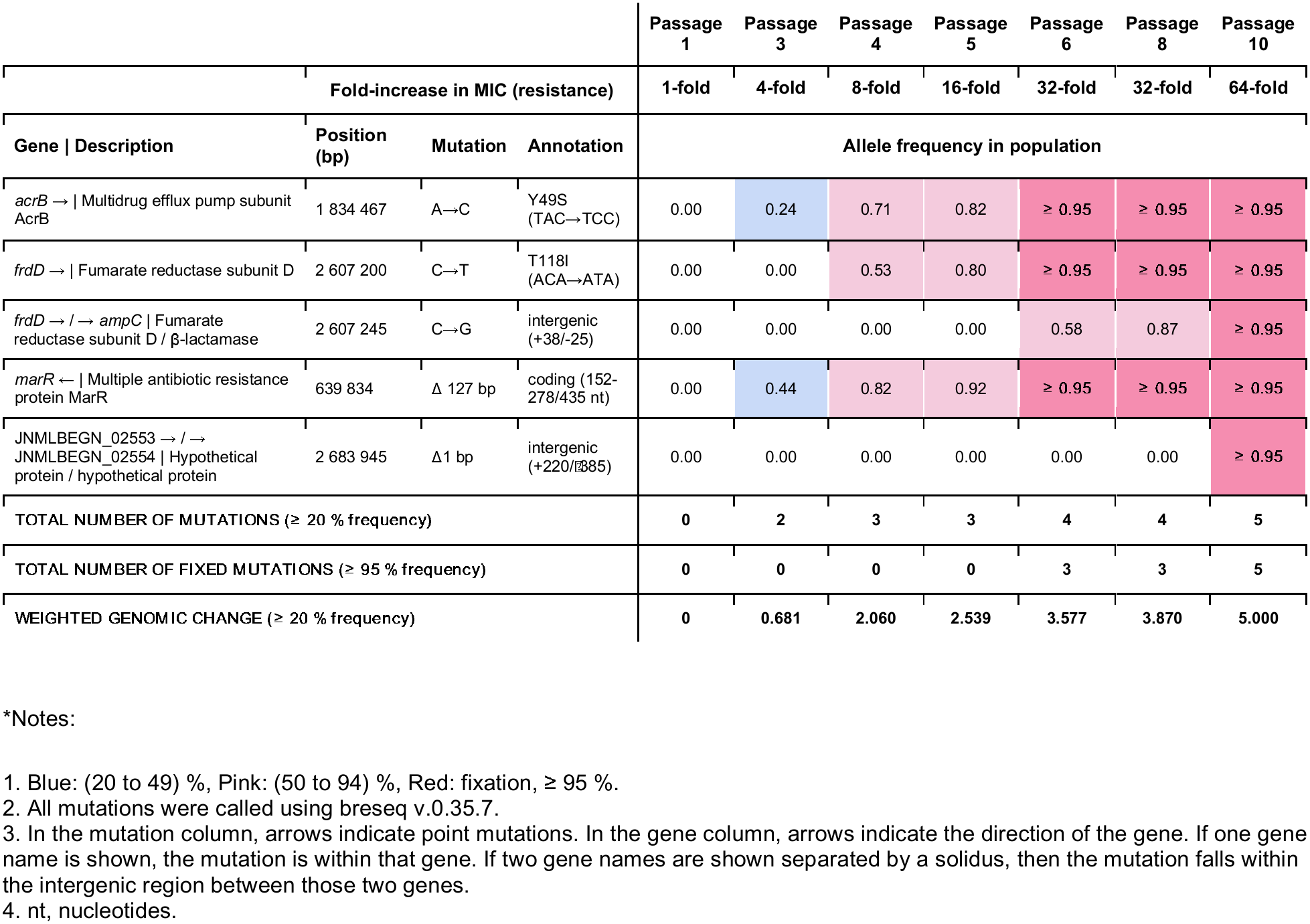
Observed mutations across passage in high R-Amp avian *E. coli* (replicate 4, Figure 1). Mutations appearing at > 40 % frequency in at least one of the populations and not present in the untreated control are included. Total number of mutations and certain mutations (and their frequency in the population) correlate with the largest increases in MIC over the 10-d evolution, meaning mutations accumulate in their effect on phenotypic level of resistance acquired. If the right mutations are picked up initially (e.g., *acrB, marR,* fumarate reductase, β-lactamase), as it appears to be the case in this lineage, resistance can build/emerge quickly and reach high levels.

| Gene \| Description | Position (bp) | Mutation | Annotation | Passage 1 | Passage 3 | Passage 4 | Passage 5 | Passage 6 | Passage 8 | Passage 10 |
|---|---|---|---|---|---|---|---|---|---|---|
|  | **Fold-increase in MIC (resistance)** |  |  | **1-fold** | **4-fold** | **8-fold** | **16-fold** | **32-fold** | **32-fold** | **64-fold** |
| **Gene \| Description** | **Position (bp)** | **Mutation** | **Annotation** | **Allele frequency in population** |  |  |  |  |  |  |
| *acrB* → \| Multidrug efflux pump subunit AcrB | 1 834 467 | A→C | Y49S<br>(TAC→TCC) | 0.00 | 0.24 | 0.71 | 0.82 | ≥ 0.95 | ≥ 0.95 | ≥ 0.95 |
| *frdD* → \| Fumarate reductase subunit D | 2 607 200 | C→T | T118I<br>(ACA→ATA) | 0.00 | 0.00 | 0.53 | 0.80 | ≥ 0.95 | ≥ 0.95 | ≥ 0.95 |
| *frdD* → / → *ampC* \| Fumarate reductase subunit D / β-lactamase | 2 607 245 | C→G | intergenic<br>(+38/-25) | 0.00 | 0.00 | 0.00 | 0.00 | 0.58 | 0.87 | ≥ 0.95 |
| *marR* ← \| Multiple antibiotic resistance protein MarR | 639 834 | Δ 127 bp | coding (152-278/435 nt) | 0.00 | 0.44 | 0.82 | 0.92 | ≥ 0.95 | ≥ 0.95 | ≥ 0.95 |
| JNMLBEGN_02553 → / → JNMLBEGN_02554 \| Hypothetical protein / hypothetical protein | 2 683 945 | Δ1 bp | intergenic<br>(+220/[illegible]385) | 0.00 | 0.00 | 0.00 | 0.00 | 0.00 | 0.00 | ≥ 0.95 |
| TOTAL NUMBER OF MUTATIONS (≥ 20 % frequency) |  |  |  | **0** | **2** | **3** | **3** | **4** | **4** | **5** |
| TOTAL NUMBER OF FIXED MUTATIONS (≥ 95 % frequency) |  |  |  | **0** | **0** | **0** | **0** | **3** | **3** | **5** |
| WEIGHTED GENOMIC CHANGE (≥ 20 % frequency) |  |  |  | **0** | **0.681** | **2.060** | **2.539** | **3.577** | **3.870** | **5.000** |

\*Notes:
1. Blue: (20 to 49) %, Pink: (50 to 94) %, Red: fixation, ≥ 95 %.
2. All mutations were called using breseq v.0.35.7.
3. In the mutation column, arrows indicate point mutations. In the gene column, arrows indicate the direction of the gene. If one gene name is shown, the mutation is within that gene. If two gene names are shown separated by a solidus, then the mutation falls within the intergenic region between those two genes.
4. nt, nucleotides.

Longitudinal analysis of the genomic amplification events further highlighted these divergent trajectories. In replicate 3A, copy number gains were detectable by day 3 and expanded markedly by day 7, with both the 149.5 kbp *acrAB*-containing region and the 243.5 kbp *ampC*-containing region showing amplification by day 10. In replicate 4A, small regions of increased copy number appeared by day 3, but amplification of the *ampC*-containing region was more gradual than in replicate 3A, becoming prominent only after day 6 and persisting through day 10, with no amplification of the *acrAB*-containing region taking place (**Figure S6**).

Taken together, the results indicate that these replicates diverged markedly in response to ampicillin pressure. Replicate 3A adopted an early fitness strategy under ampicillin, improving tolerance even without attendant MIC gains and allowing populations to persist until a conventional resistance trajectory (e.g., *acrB, frdD/ampC, marR*) emerged. Replicate 4A, by contrast, rapidly acquired and fixed canonical AST-defined determinants. Despite these distinct adaptive trajectories, both lineages exhibited motility losses with broadly similar timing and magnitude. In other words, multiple distinct resistance states can produce similar motility phenotypes. Notably, motility did not scale proportionally (neither directly nor consistently) with MIC across serial passages, indicating a non-bijective, temporally asynchronous relationship between resistance level and swimming behavior, which makes sense as the evolutionary processes that govern fitness are complex and multidimensional. Accordingly, motility loss may be better interpreted as a conserved adaptive response to antibiotic stress than as a direct consequence of specific resistance mutations or amplifications. In some contexts (e.g., replicate 3A), motility decline preceded or exceeded resistance measured by MIC; in others (e.g., replicate 4A), motility lagged resistance acquisition or varied independently. The most parsimonious explanation is pleiotropy in proton motive force (PMF) linked physiology: both efflux activity and flagellar rotation draw on this energy gradient, so early perturbations to PMF dependent processes can manifest as motility defects before high level resistance is phenotypically evident. Furthermore, resistance may have become constitutively expressed through the regulatory mutations in global regulator MarR, resulting in persistent activation of resistance pathways (sustained efflux upregulation) even in the absence of antibiotics and potentially contributing to fitness and motility tradeoffs; motility loss may, in turn, mitigate the costs associated with constitutive resistance. With respect to long-term maintenance of resistance, an important unresolved question — and a focus of future investigation — is whether compensatory evolution can progressively restore aspects of fitness, potentially including motility, depending on the genetic basis of the compensatory mutations.

These insights have broader implications for how antibiotic resistance emergence is detected and interpreted, motivating consideration of phenotype-based metrics as complementary tools for AMR surveillance. Conventional genetic and clinical AST measurements fail to capture early physiological adaptations to antibiotic stress. Yet, as we have seen in this work, these responses are critical in improving the growth, survival, and recovery of bacterial populations under antibiotic exposure and ultimately determining later resistance phenotypes. Changes in bacterial motility, particularly reversible reductions linked to PMF-dependent efflux, thus may bode well to serve as an early phenotype-level indicator of emergent efflux-based β-lactam resistance. Because motility reflects integrated cellular energetics rather than individual genetic determinants, motility-based readouts could, in principle, provide faster or more scalable detection of certain resistance strategies than gene-specific assays such as PCR targeting *acrB* or *marR*. Approaches leveraging microscopy or microfluidic platforms to quantify swimming behavior may therefore enable rapid screening for efflux-associated AMR phenotypes without prior knowledge of the underlying genotype. Such platforms could have applications in environmental surveillance (*e.g.,* water or food testing), clinical diagnostics, and microbiome monitoring, where rapid determination of the presence, absence, or abundance of resistant bacteria is critical. While further validation is needed, our findings highlight motility as a potentially informative and underutilized phenotypic readout for AMR detection and risk assessment.

## 4. Conclusion

Our results emphasize that adaptive evolution extends beyond canonical, AST-defined resistance mechanisms and includes broader physiological reprogramming that shapes tolerance and persistence. Specifically, our findings suggest that flagellar motility is among the earliest cellular functions to be downregulated during the evolution of β-lactam resistance. Prolonged antibiotic exposure consistently selected for loss of flagellar motility, demonstrating a clear association between reduced swimming activity and increased fitness under β-lactam stress. Although resistance trajectories diverged markedly across replicates, motility loss converged, highlighting motility as a broadly selected adaptive trait under antibiotic stress rather than a marker of a single resistance mechanism. Rather than representing independent resistance determinants, these adaptations may reflect an epistatic resistance network, in which *marR* deletion reshapes the adaptive landscape and changes the phenotypic consequences of mutations in efflux loci such as *acrB*, while favoring physiological tradeoffs such as motility loss. The mutations identified across evolved populations point to coordinated shifts in metabolism, membrane energetics, and efflux capacity. These systems-level adaptations likely reshape cellular resource allocation, favoring survival under antibiotic pressure at the expense of energetically costly processes such as flagellar biosynthesis and rotation. Importantly, these early physiological adaptations, though not captured by standard resistance metrics, significantly enhance growth, survival, and recovery under antibiotic exposure. This work thus motivates the development of phenotypic separation or enrichment strategies that exploit motility differences to detect or isolate antimicrobial-resistant subpopulations.

Microfluidic or motility-based sorting platforms could be designed to enrich non-motile or weakly motile cells, which may be disproportionately associated with energetically costly, efflux-driven resistance states. By integrating phenotypic screening with genetic and MIC-based approaches, future work may better resolve early adaptive states that precede — and potentially enable — the emergence of clinically defined resistance.

## DISCLAIMER

Certain commercial equipment, instruments, software, or materials are identified in this paper in order to specify the experimental procedure adequately. Such identification is not intended to imply recommendation or endorsement by the National Institute of Standards and Technology, nor is it intended to imply that the materials, software, or equipment identified are necessarily the best available for the purpose.

The National Institute of Standards and Technology Research Protections Office determined this project is “not human subjects research.”

Official contribution of the National Institute of Standards and Technology; not subject to copyright in the United States.

