## Supplemental Information for "Reversible decrease in swimming motility accompanies emergent β-lactam antibiotic resistance in *Escherichia coli*"

**Table of Contents:**

**Supplementary Fig. 1** – Effect of Temperature and Time on Motility of Untreated *E. coli*….P. S2

**Supplementary Fig. 2** – Swimming Motility in Selective Soft Agar…………………………....P. S2

**Supplementary Fig. 3** – Swimming Motility of Cultures Grown in Non-Selective Media……P. S3

**Table S1** – Baseline Antibiotic Susceptibility and Cross-Resistance Patterns…..……………P. S4 **Table S2** – Mann-Whitney Analysis of Swim Zone Halo Diameters……………………………P. S4 **Table S3** – Dunn’s Post Hoc Analysis of Ampicillin Clone Replicates.……………...…………P. S5

**Table S4** – Pairwise Comparisons of Ampicillin Replicate Populations.……………………….P. S5

**Table S5** – Pairwise Comparisons of Cephalexin Replicate Populations…………………......P. S5

**Table S6** – Pairwise Comparisons of Untreated Replicate Populations……………………….P. S6

**Supplementary Fig. 4** – Motility Response Following Removal of Antibiotic Pressure…......P. S6

**Supplementary Fig. 5** – Mutation Frequency Following Removal of Antibiotic Pressure…...P. S7 **Table S7** – Mutation Frequency Following Removal of Antibiotic Pressure…………………...P. S8 **Table S8** – Genetic Basis of Baseline Resistance in the Ancestral Strain………………..……P. S9

**Supplementary Fig. 6** – Copy Number Variation Analysis Showing Amplification …….......P. S10 **Supplementary Fig. 7** – Detailed Time-Course Genomics Analysis for Replicate 3A..........P. S11

**References**…………………………………………………………………………………………..P. S12

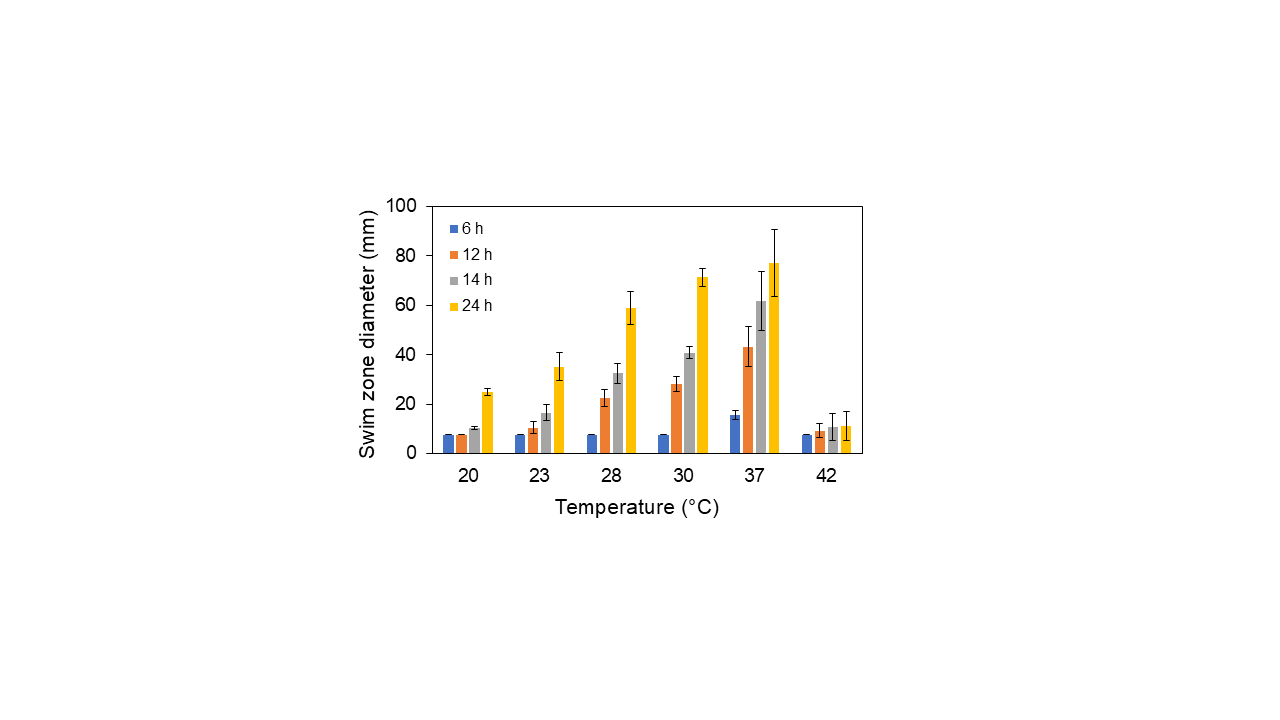

**Supplementary Fig. 1. Temperature- and time-dependent motility results for untreated *E. coli*.** Error bars represent ± 1 standard deviation of the mean for *n = 3* independent biological replicates.

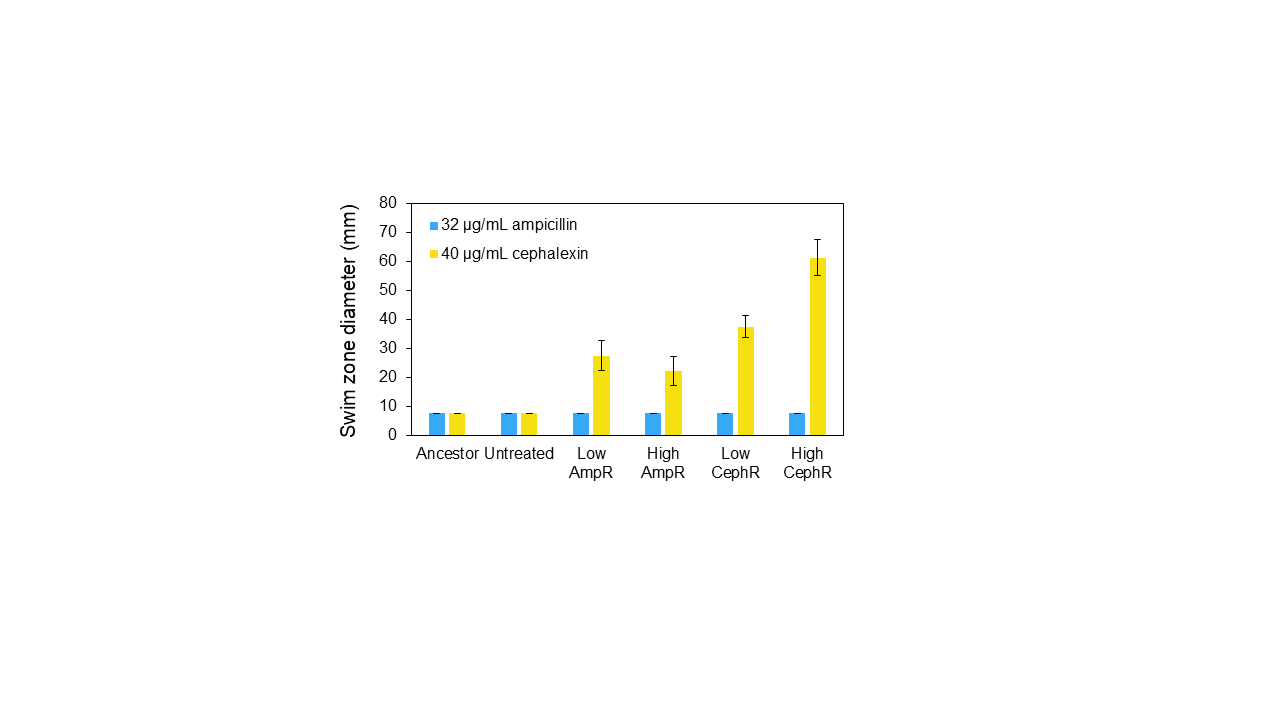

**Supplementary Fig. 2. Swimming Motility in Selective Soft Agar.** Swimming motility in soft agar supplemented with cephalexin (blue) or ampicillin (yellow-orange), illustrating differential motility responses under antibiotic exposure and highlighting antibiotic-specific effects on motility. Note: Low → Intermediate (I-Amp; MIC = 16 μg/mL); High → Resistant (R-Amp; MIC ≥ 32 μg/mL). Error bars represent ± 1 standard deviation of the mean for *n = 3* independent biological replicates.

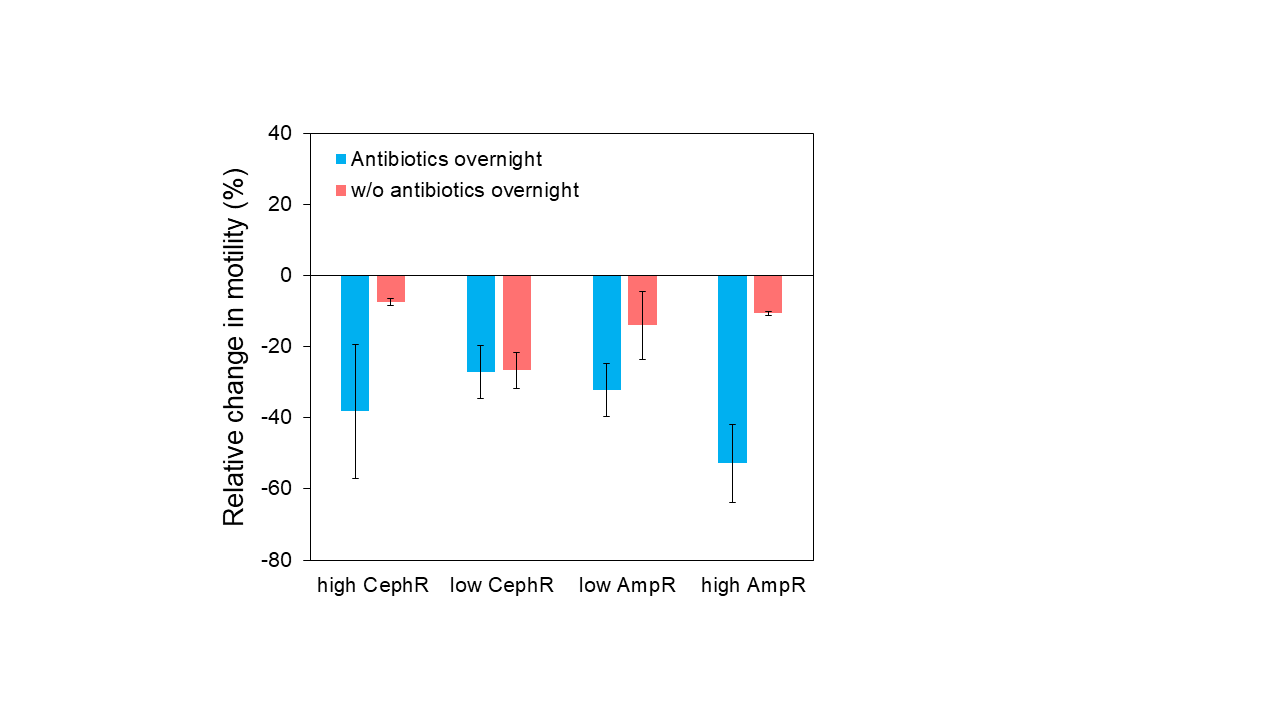

**Supplementary Fig. 3. Swimming Motility of Cultures Grown in Non-Selective Media.** Motility characterization following different overnight culture methods. Swimming motility of AmpR and CephR *E. coli* populations after 10 passages was assessed following revival from frozen stocks (–80 °C) and overnight growth in either antibiotic-containing or non-selective media. When antibiotics were omitted from the overnight culture, motility reduction was less pronounced. This likely reflects relaxation of selection, allowing more motile, antibiotic-sensitive subpopulations to persist or outcompete highly resistant, reduced-motility strains. In other words, cultures grown in the absence of antibiotics showed more motile behavior, likely as a result of less resistant phenotypes present in this mixed population. When the pressure is removed, even after one passage in non-selective media, motility recovers, highlighting the role of the overnight culture conditions and continuing/maintaining the selection in preserving the reduced-motility phenotype (and/or selecting for the resistant subpopulations). Thus, motility reduction appears to represent a transient stress response to sublethal antibiotic exposure and an unstable phenotype, even when associated with high-level resistance evolution under strong selection. Note: Low → Intermediate (I-Amp or I-Ceph; MIC = 16 μg/mL); High → Resistant (R-Amp or R-Ceph; MIC ≥ 32 μg/mL). Error bars represent ± 1 standard deviation of the mean for *n = 3* independent biological replicates.

**Table S1.** Changes in antibiotic susceptibility and emergence of cross-resistance in *E. coli* mixed populations. Baseline resistance and cross-resistance was assessed in the ancestral strain as well as the lab-evolved intermediate (I-Amp, I-Ceph) and resistant (R-Amp, R-Ceph) populations after 10 d of antibiotic selection using the Thermo Scientific Sensititre Gram Negative (GN4F) Plate with FDA interpretive criteria for Enterobacteriaceae. Assays were run in triplicate. Lab-evolved resistant strains exhibited cross-resistance to multiple antibiotics, including those within similar classes (e.g., penicillins, first- and third-generation cephalosporins, β-lactamase inhibitors) as well as across different classes (e.g., tetracyclines).

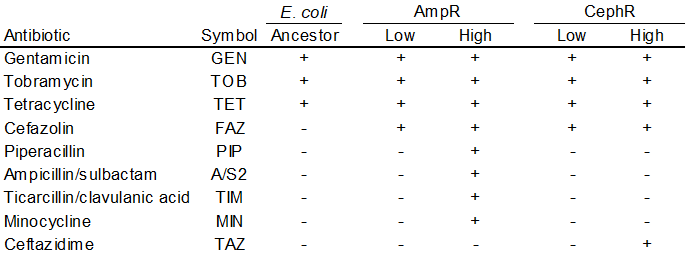

*Note: Low → suceptible/Intermediate (I-Amp; MIC < 8 μg/mL); High → Resistant (R-Amp; MIC ≥ 32 μg/mL)

**Table S2.** Summary of additional statistical analyses (Mann-Whitney) performed on raw swimming motility data collected on day 10, grouped by treatment. Swimming motility, measured by swim halo diameter, was significantly reduced in antibiotic-exposed populations compared with untreated populations.

|  | Populations | | Clones |
| --- | --- | --- | --- |
|  | Ampicillin | Cephalexin | Ampicillin |
| Untreated | *p* = 0.024 | *p* = 0.024 | *p* = 0.075 |
| Ampicillin | -- | *p* = 0.841 | -- |

**Table S3.** Dunn’s post hoc (Kruskal-Wallis multiple comparison) analysis of swim halo diameter compared between individual *clones* exposed to ampicillin. P-values were adjusted with the Bonferroni method.

|  | Comparison | Z | P.unadj | P.adj |
| --- | --- | --- | --- | --- |
| 1 | A1 vs A2 | 0.15 | 0.88 | 1.00 |
| 2 | A1 vs A3 | 1.19 | 0.23 | 1.00 |
| 3 | A2 vs A3 | 0.88 | 0.38 | 1.00 |
| 4 | A1 vs A4 | 3.38 | 0.00072 | 0.0072 |
| 5 | A2 vs A4 | 2.74 | 0.0061 | 0.061 |
| 6 | A3 vs A4 | 2.50 | 0.012 | 0.12 |
| 7 | A1 vs A5 | 3.30 | 0.00097 | 0.0096 |
| 8 | A2 vs A5 | 2.69 | 0.0071 | 0.071 |
| 9 | A3 vs A5 | 2.42 | 0.016 | 0.16 |
| 10 | A4 vs A5 | 0.00 | 1.00 | 1.00 |

**Table S4.** Dunn’s post hoc (Kruskal-Wallis multiple comparison) analysis of swim halo diameter compared between *populations* exposed to ampicillin. P-values adjustment: Bonferroni method.

|  | Comparison | Z | P.unadj | P.adj |
| --- | --- | --- | --- | --- |
| 1 | A1 vs A2 | 0.077 | 0.94 | 1.00 |
| 2 | A1 vs A3 | 0.051 | 0.96 | 1.00 |
| 3 | A2 vs A3 | -0.031 | 0.98 | 1.00 |
| 4 | A1 vs A4 | 2.45 | 0.014 | 0.14 |
| 5 | A2 vs A4 | 2.36 | 0.018 | 0.18 |
| 6 | A3 vs A4 | 2.61 | 0.0091 | 0.091 |
| 7 | A1 vs A5 | 1.49 | 0.13 | 1.00 |
| 8 | A2 vs A5 | 1.42 | 0.16 | 1.00 |
| 9 | A3 vs A5 | 1.55 | 0.12 | 1.00 |
| 10 | A4 vs A5 | -0.78 | 0.44 | 1.00 |

**Table S5.** Dunn’s post hoc (Kruskal-Wallis multiple comparison) analysis of swim halo diameter compared between populations exposed to cephalexin. P-value adjustment: Bonferroni method.

|  | Comparison | Z | P.unadj | P.adj |
| --- | --- | --- | --- | --- |
| 1 | C1 vs C2 | -0.49 | 0.62 | 1.00 |
| 2 | C1 vs C3 | -1.42 | 0.16 | 1.00 |
| 3 | C2 vs C3 | -0.67 | 0.51 | 1.00 |
| 4 | C1 vs C4 | 1.45 | 0.15 | 1.00 |
| 5 | C2 vs C4 | 1.68 | 0.093 | 0.93 |
| 6 | C3 vs C4 | 2.61 | 0.0092 | 0.092 |
| 7 | C1 vs C5 | -0.23 | 0.82 | 1.00 |
| 8 | C2 vs C5 | 0.23 | 0.82 | 1.00 |
| 9 | C3 vs C5 | 0.93 | 0.35 | 1.00 |
| 10 | C4 vs C5 | -1.45 | 0.15 | 1.00 |

**Table S6.** Dunn’s post hoc (Kruskal-Wallis multiple comparison) analysis of swim halo diameter compared across untreated populations. P-values were adjusted with the Bonferroni method.

|  | Comparison | Z | P.unadj | P.adj |
| --- | --- | --- | --- | --- |
| 1 | U1 vs U2 | 1.78 | 0.075 | 0.75 |
| 2 | U1 vs U3 | -0.23 | 0.82 | 1.00 |
| 3 | U2 vs U3 | -2.01 | 0.044 | 0.44 |
| 4 | U1 vs U4 | 1.19 | 0.23 | 1.00 |
| 5 | U2 vs U4 | -0.59 | 0.55 | 1.00 |
| 6 | U3 vs U4 | 1.42 | 0.16 | 1.00 |
| 7 | U1 vs U5 | 2.06 | 0.040 | 0.40 |
| 8 | U2 vs U5 | 0.27 | 0.78 | 1.00 |
| 9 | U3 vs U5 | 2.29 | 0.022 | 0.22 |
| 10 | U4 vs U5 | 0.87 | 0.38 | 1.00 |

Pairwise comparisons among the five replicate clones revealed that replicates A4 and A5 had significantly higher motility than replicate A1 (Bonferroni-adjusted *p* = (0.007 to 0.01)), whereas no other pairwise differences were statistically significant. Notably, in the unadjusted tests, replicates A4 and A5 also differed from replicates A2 and A3 (*p* < 0.05), although these differences were not significant after correction for multiple comparisons.

For replicate populations exposed to ampicillin and cephalexin, as well as untreated conditions, no pairwise differences were statistically significant after correcting for multiple comparisons, indicating that motility was generally consistent across replicates within each condition.

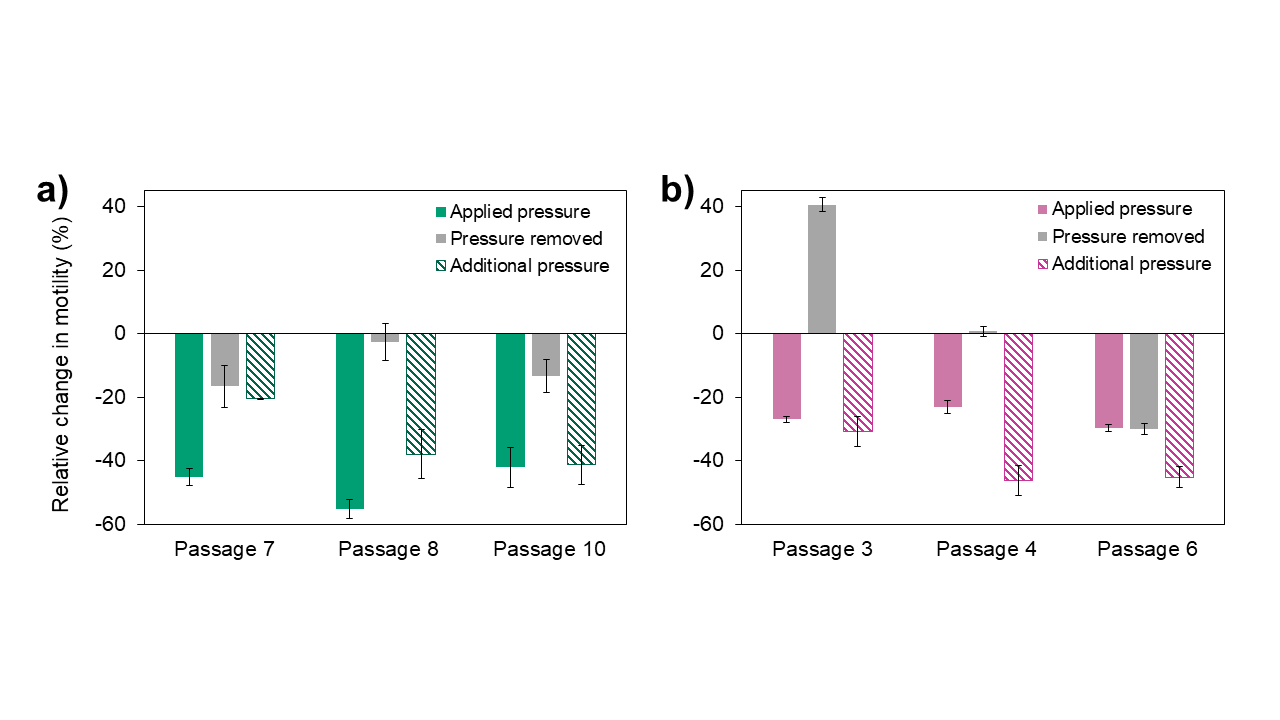

**Supplementary Fig. 4**. **Motility Response Following Removal of Antibiotic Pressure.** Swimming motility of *E. coli* populations from (a) replicate 3A (I-Amp) and (b) replicate 4A (R-Amp) measured at selected passages along their evolutionary trajectories relative to the wild type. Time points were chosen based on the population frequencies of *marR* and/or *acrB* mutations. Motility was assessed after removal of antibiotic selection for three passages and after re-exposure for an additional three passages. In most cases, motility returned to approximately wild-type levels following removal of selection, whereas renewed antibiotic exposure produced comparable reductions in swimming motility. Error bars represent ± 1 standard deviation of the mean for *n = 3* independent biological replicates.

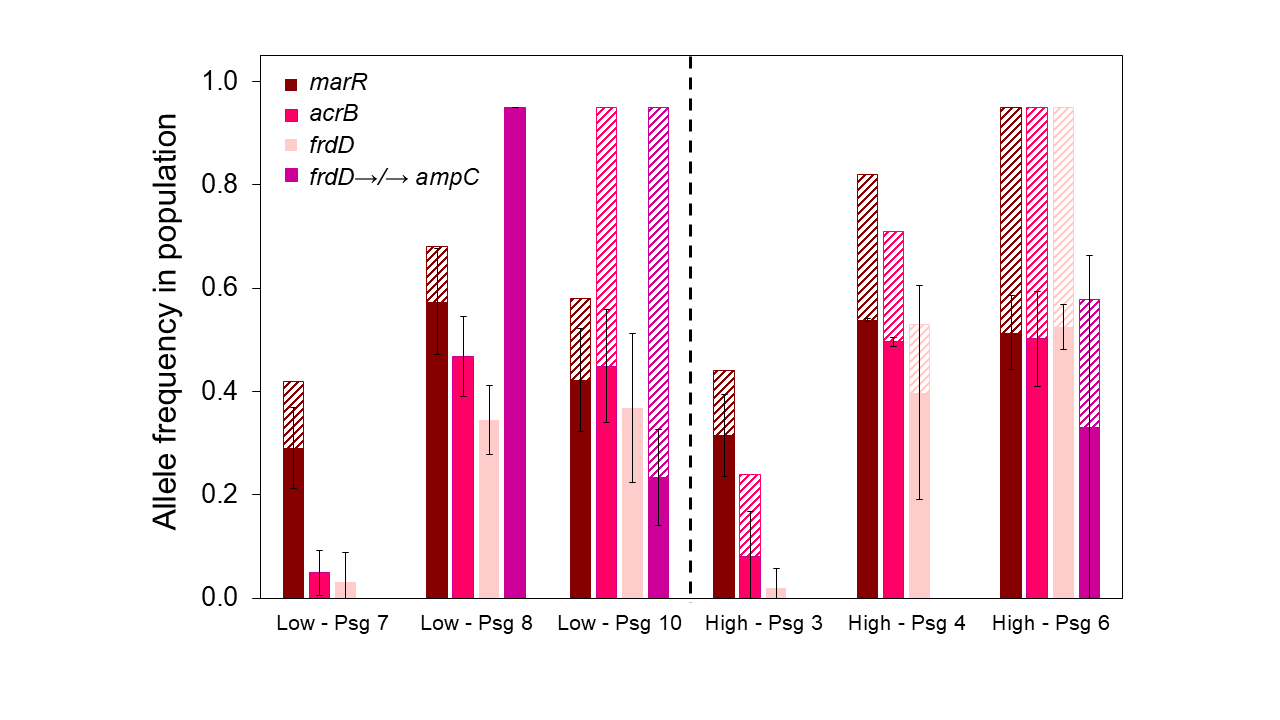

**Supplementary Fig. 5. Mutation Frequency Following Removal of Antibiotic Pressure.** Percent frequency of major mutations following removal of selective pressure at the identified passage number for three consecutive passages. Dashed bars represent mutation frequencies prior to pressure removal, and solid bars represent frequencies after removal. In most cases, mutation frequencies decrease following release from selection. Error bars represent ± 1 standard deviation of the mean for *n = 3* independent biological replicates per condition after pressure removal.

**Table S7**. Mutation frequency following removal of antibiotic pressure. Level of stability of observed mutations remaining after antibiotic pressure is removed across select passages in replicate 3A (I-Amp) and replicate 4A (R-Amp). Mutations exceeding a frequency of 20 % in at least one replicate population and not present in the untreated control are included. Values represent the average of *n* = 3 replicates sequenced for each condition, with the numbers in parentheses representing the standard deviation. It is important to note that no additional mutations exceeding 20 % frequency arose in genes other than those already identified under selective pressure. In most cases, motility was restored to wild-type levels after removal of the selective pressure. Concurrently, the frequency of these mutations declined within the population, supporting their role in driving the reduced motility phenotype. Together, these findings further corroborate a causal link between antibiotic pressure and mutations underlying motility loss.

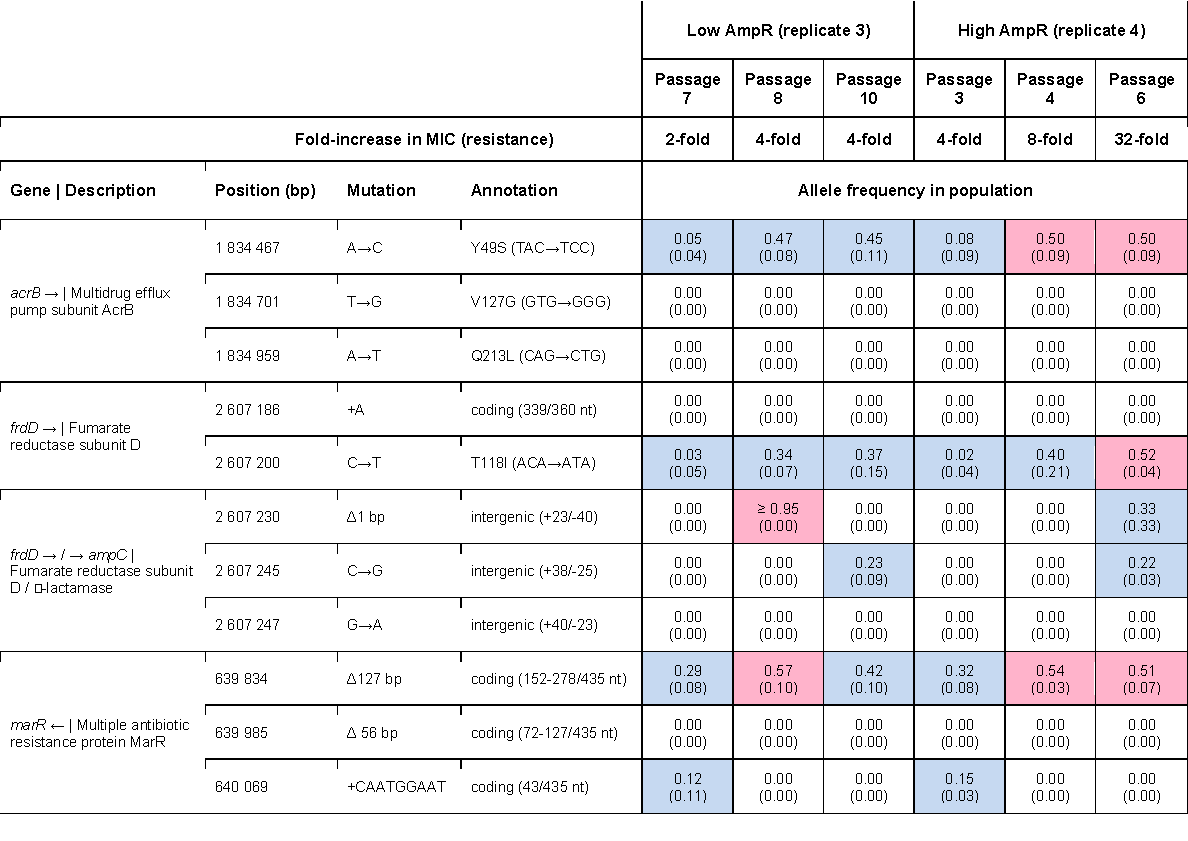

**Table S8**. Genetic basis of baseline resistance in the ancestral strain. The presence of antimicrobial resistance and virulence genes was determined using AMRFinderPlus v4.2.5.^1^

|  | Size (bp) | Description | Antibiotic Resistance Genes | Virulence-Associated Genes |
| --- | --- | --- | --- | --- |
| Chromosome | 4 793 125 | *E. coli* NIST 0056 chromosome | - *ampC*: chromosomal class C β‑lactamase - *tetA*/*tetR*: tetracycline resistance with repressor - *ant1*: streptomycin resistance - *aphA*, *yokD*: aminoglycoside resistance - *abaF*: fosfomycin resistance - *bcr*: bicyclomycin resistance - *macA*/*B*: macrolide resistance - *acrA*, *acrB*, *acrE*/*F*/*Z*, *mdfA*, *emrA*/*B*/*E*/*K*/*Y*, *mdtA*-*O* (subset) - MDR efflux (RND/MFS/ABC) - *marA*/*R,* *soxR*/*S*, *rob*, *baeS*/*R*: MDR regulators - *sbmA*: peptide antibiotic transporter | - long polar fimbriae (*lpfA*) - ETEC adhesins (*cfaB/E*) - UPEC adhesins (*papC/D)* - T3SS (s*paO/P/Q/R/S, prgH, orgA, sicA*) - T6SS (*hcp1*, *vgrG1*, *clpV1*, *vapB/C*) - toxins/effectors (*hlyE, cdiA, sopB*, *shlB*, *vapC*) - aerobactin siderophore (*iucABCD–iutA*) |
| Plasmid 1 | 122 629 | IncI1-type conjugative plasmid with multiple resistance and virulence determinants | - *aac(3)-VIa*: aminoglycoside resistance - *aadA1*: streptomycin resistance (pseudogene) - *sul1*: sulfonamide resistance - *tetA*/*tetR*: tetracycline resistance with repressor - *qacEΔ1*: quaternary ammonium compound efflux | - conjugation system (*tra*/*trb*) - type IV pilus biogenesis (*pil* genes) - bacteriocin (colicin-related genes) - mercury resistance (*mer* operon) - toxin-antitoxin systems (*yacA*/*B*, *hok*/*gef* family) |
| Plasmid 2 | 148 536 | ColV-like conjugative and multifunctional virulence plasmid, similar to APEC virulence plasmids | - *mig-14*: antimicrobial resistance - efflux-related genes (*macB*, *tolC*, *crcB*) | - conjugation system - siderophore systems (aerobactin operon, salmochelin operon, *sitABCD* system) - serum survival (*iss*) - complement resistance (*traT*) - toxin-antitoxin systems (*vapB*/*C*, *yacA*/*B*, *relE, phd*/*yefM*, *hok*) - bacteriocin (colicin-related genes) |
| Plasmid 3 | 111 610 | Cryptic plasmid with replication and stability functions; contains multiple overlapping/duplicated replication regions suggesting recombination events | - none detected | - toxin-antitoxin systems (*yafQ*, *relB/dinJ*, *phd*/*yefM*) |
| Plasmid 4 | 93 185 | Cryptic prophage plasmid with extensive lysogenic phage integration machinery | - *bmh*: bleomycin hydrolase - *terB*: tellurite resistance - multiple efflux and DNA protection systems (N-6 DNA methylase) | - multiple phage-related genes (phage structural genes, phage regulatory and lysogeny genes) - toxin-antitoxin systems (*phd*/*yefM, doc*) |

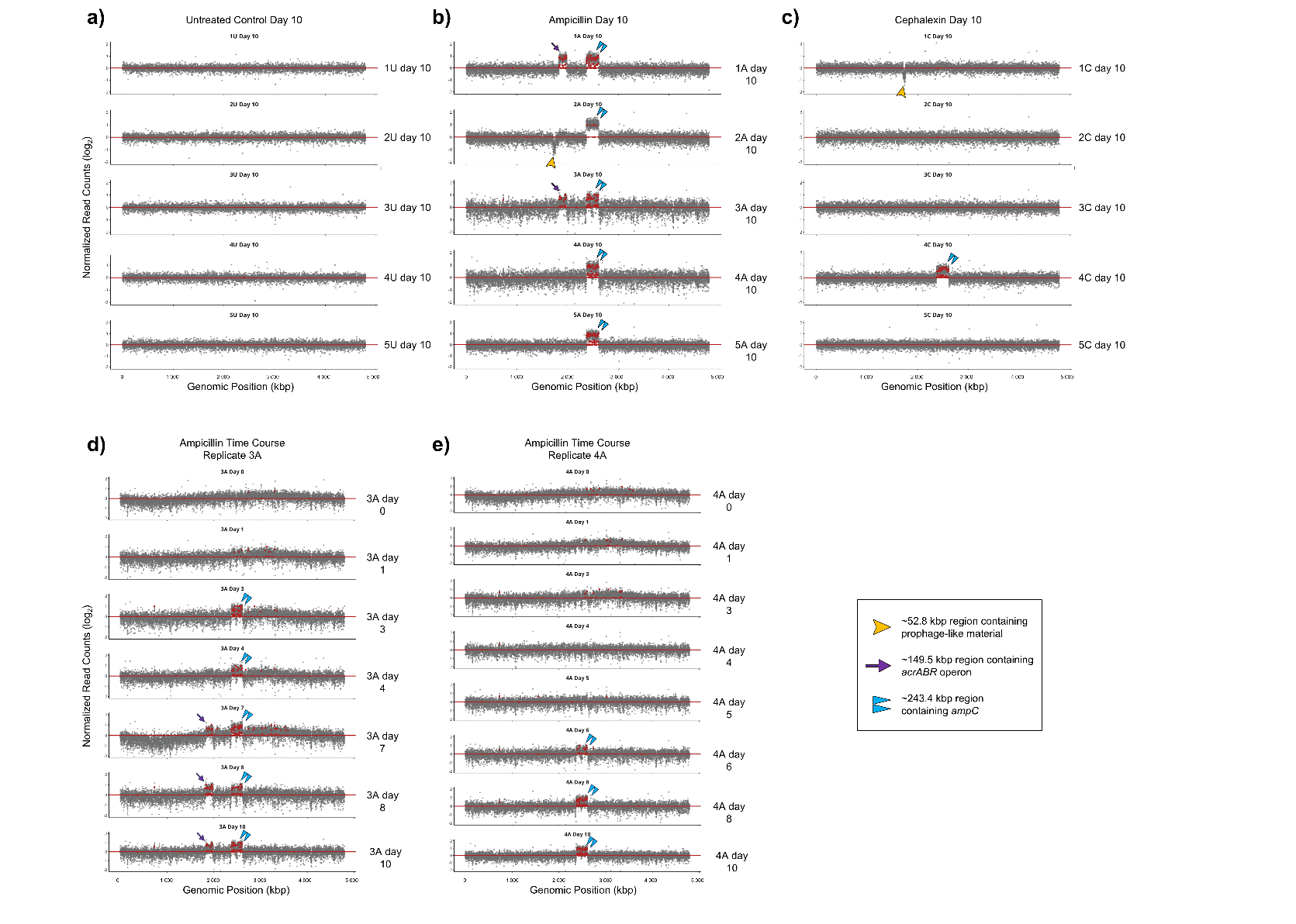

**Supplementary Fig. 6. Copy Number Variation Analysis Showing Genomic Amplifications.** Normalized sequencing coverage across the indicated region for each replicate reveals duplicated segments in NIST 0056 under (a) no selection; (b, d, e) ampicillin; and (c) cephalexin. Panels (b, c) show five replicates at day 10, whereas (d, e) display time-series data for replicates 3A and 4A, respectively. Single purple arrows indicate regions containing *acrA* (and other genes within the same operon), double blue arrows denote regions containing *ampC*, and yellow arrows mark prophage-like elements.

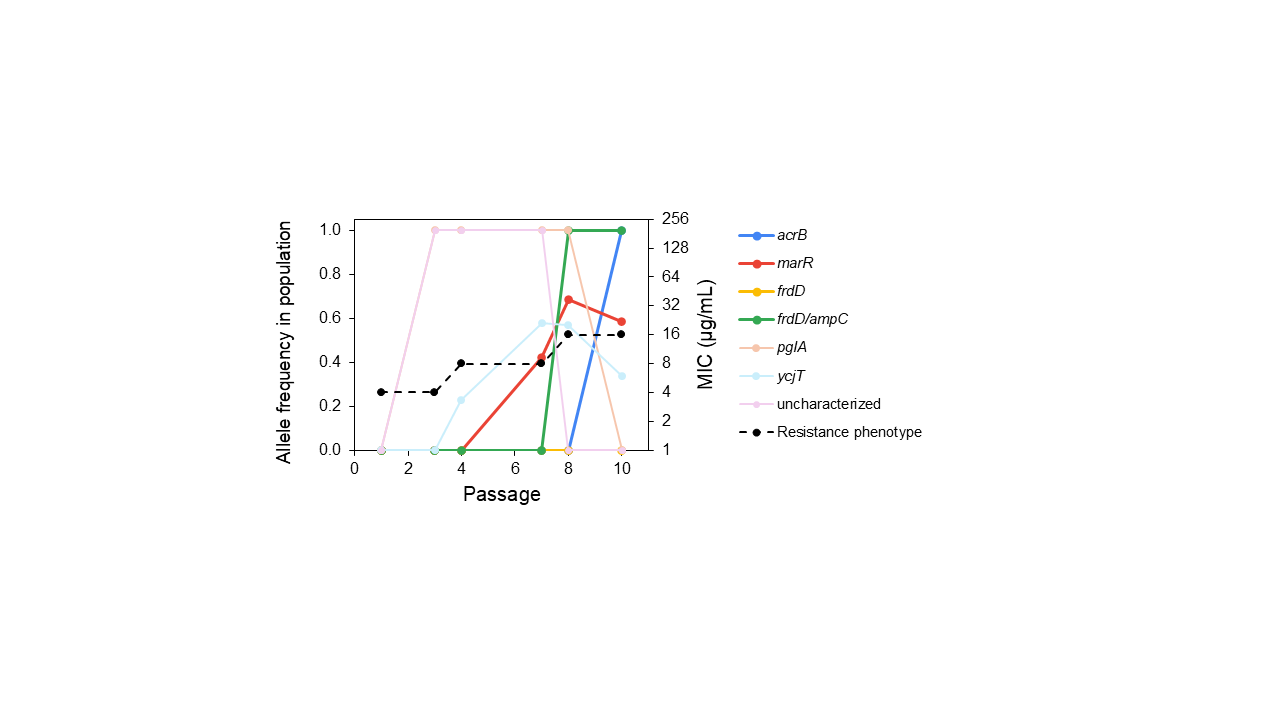

**Supplementary Fig. 7. Detailed Time-Course Genomics Analysis for Replicate 3A.** Mutations in non-canonical AMR-associated genes, (*pglA, ycjT*, and an uncharacterized protein) rose to fixation by day 3 and were maintained at 100 % frequency until days 7 to 8, after which they were completely lost from the population.
